# Deciphering the molecular mechanism underlying morphology transition in two-component DNA-protein cophase separation

**DOI:** 10.1101/2024.01.26.577305

**Authors:** Cheng Li, Yunqiang Bian, Yiting Tang, Lingyu Meng, Peipei Yin, Ye Hong, Jun Cheng, Yuchen Li, Jie Lin, Chao Tang, Chunlai Chen, Wenfei Li, Zhi Qi

**Affiliations:** Center for Quantitative Biology, Peking-Tsinghua Center for Life Sciences, Academy for Advanced Interdisciplinary Studies, Peking University, Beijing 100871, China; Wenzhou Key Laboratory of Biophysics, Wenzhou Institute, University of Chinese Academy of Sciences, Wenzhou, Zhejiang 325000, China; School of Life Sciences, Beijing Advanced Innovation Center for Structural Biology, Beijing Frontier Research Center for Biological Structure, State Key Laboratory of Membrane Biology, Tsinghua University, 100084 Beijing, China; The Integrated Science Program, Yuanpei College, Peking University, Beijing 100871, China; School of Physics, Peking University, Beijing 100871, China; Department of Physics, National Laboratory of Solid-State Microstructure, and Collaborative Innovation Center of Advanced Microstructures, Nanjing University, Nanjing 210093, China

## Abstract

Nucleic acid and protein co-condensates exhibit diverse morphologies crucial for fundamental cellular processes. Despite their significance, the molecular mechanisms governing morphology transitions remain poorly understood. To address this gap in knowledge, we investigated DNA and the human transcription factor p53 as a model system, specifically focusing on DNA-protein interactive co-condensates (DPICs)—a scenario where neither dsDNA nor the protein demonstrates phase-separation behavior individually. Through a combination of experimental assays and theoretical approaches, we elucidated: (i) the phase diagram of DPICs, identifying two distinct transition phenomena—a phase transition between viscoelastic fluid and viscoelastic solid states, and a morphology transition from droplet-like to "pearl chain"-like DPICs; (ii) the growth dynamics of DPICs. Droplet-like and "pearl chain"-like DPICs, although with dramatically distinct final morphologies and material properties, share a common initial critical microscopic cluster (CMC) size at the nanometer scale during the early stage of phase separation. These findings provide novel insights into the biophysical mechanisms underlying multi-component phase separations within cellular environments.

**Significance Statement:** Nucleic acids and proteins have the capacity to form co-condensates, exhibiting various morphologies, including droplet-like and “pearl chains” formations. Despite this observation, the underlying biophysical mechanisms remain poorly understood. In this study, we employed DNA and the protein p53 as a model system. Our investigation revealed that the strength of the DNA-p53 interactions dictates the material properties of the co-condensates, leading to a transition from a viscoelastic fluid to a viscoelastic solid phase. This transition is accompanied by a morphological shift from droplet-like formations to structures resembling “pearl chains”. Additionally, we explored the growth dynamics of these co-condensates and demonstrated that the strength of p53-DNA interactions influences the relaxation time of the co-condensates, thereby potentially determining their morphological features.

## Introduction

Eukaryotic cells utilize lipid membranes to compartmentalize intracellular organization. However, recent advancements in *in vivo* and *in vitro* research have unveiled an alternative mechanism employed by these cells to organize their complex biochemistry (1). Specifically, macromolecules, such as nucleic acids and proteins, have been identified to undergo liquid-liquid phase separation (LLPS), leading to the assembly of membrane-lacking compartments (2-6). Two primary driving forces for the formation of LLPS include protein intrinsically disordered regions (IDRs) (7) and/or multivalent interactions among modular biomacromolecules (8). These membrane-lacking compartments are recognized as biomolecular condensates, playing pivotal roles in various biological functions. Importantly, the dysregulation of condensation processes has been implicated in a spectrum of human diseases (4).

Biomolecular condensates, arising from individual or multiple biomacromolecular species, can exist as either single-component or multi-component entities. Notably, nucleic acids have emerged as pivotal scaffolds in orchestrating the formation of nucleic acid-protein co-condensates (NAPCs), a distinct subclass of two-component condensates (9). NAPCs encompass a spectrum of assemblies, including RNA-protein co-condensates (10-15), single-stranded DNA (ssDNA)-protein co-condensates (16), and double-stranded DNA (dsDNA)-protein co-condensates (17-29).

During the typical process of LLPS, liquid-like condensates often fuse together, leading to the formation of larger spherical compartments with higher circularity values. Interestingly, NAPCs have been observed to exhibit “pearl chain”-like morphologies characterized by lower circularity values, which usually behave as gel/solid-like characteristics. For instance, nucleosomal arrays, a type of dsDNA-protein co-condensates, have been reported to adopt both droplet-like morphologies (17, 21, 22) and “pearl chain”-like morphologies (23, 24). Similarly, various examples of RNA-protein co-condensates have been documented to display diverse condensate morphologies (10, 14, 30). A comprehensive understanding of the molecular mechanisms governing the morphologies of NAPCs is essential for elucidating fundamental cellular processes.

Previous studies (10, 14, 30) have extensively investigated the morphologies of RNA-protein co-condensates. These studies have demonstrated that enhanced intermolecular interaction strengths within RNA-protein co-condensates result in prolonged relaxation times, leading to a notable morphological transition from droplet-like RNA-protein co-condensates to “pearl chain”-like structures, accompanied by a change in material properties. However, it remains uncertain whether analogous phenomena occur in other types of NAPCs. Furthermore, the fundamental biophysical characteristics governing condensate morphology in two-component nucleic acid-protein cophase separation remain elusive, leaving critical inquiries unanswered. To shed light on the molecular mechanisms underlying morphology transition, it is imperative to address two key questions: (i) What is the relationship between material property transition and morphological transition when changing parameters during cophase separation? Answering this question necessitates the construction of a phase diagram for two-component nucleic acid-protein cophase separation; (ii) How do intermolecular interaction strengths at the nanometer scale influence the morphological transition of condensates at the micrometer scale? Addressing this question requires an exploration of the growth dynamics of NAPCs. To address these uncertainties and elucidate the physical parameters dictating NAPC morphologies, we adopt a comprehensive approach that integrates experimental and theoretical methodologies. Our investigation focuses on the simplest form of NAPCs, wherein neither dsDNA nor protein independently undergoes phase separation; rather, co-condensates emerge only upon combining both components. We refer to this model system as dsDNA-protein interactive co-condensates (DPICs).

In this study, we employed the human transcription factor p53 (31) along with dsDNA to initiate the formation of DPICs. By utilizing molecular dynamics (MD) simulations, we investigated the mechanism underlying DPIC formation, highlighting the pivotal role of p53 in bridging different DNA molecules. We demonstrated that increasing either the number of bridges on each DNA or the binding affinity between DNA and protein enhances the inter-molecular interaction strengths within DPICs, inducing a morphological transition from a droplet-like to a “pearl chain”-like structure. The viscoelastic properties of droplet-like DPICs were validated through atomic force microscopy-based force spectroscopy (AFM-FS). Moreover, we systematically mapped out the phase diagram of DPICs, highlighting two distinct transitions: a transition in physical properties and a shift in morphology. Our investigation also encompassed the utilization of dual-color fluorescence cross-correlation spectroscopy (dcFCCS) assays. These assays unveiled that droplet-like and “pearl chain”-like DPICs, although with dramatically distinct final morphologies and material properties, share a common initial critical microscopic cluster (CMC) size (radii of approximately 15-30 nm) at the nanometer scale at the early stage of phase separation. Notably, these CMCs exhibit varying viscoelastic properties and can coalesce into larger DPICs with differing morphologies and material properties.

## Results

### p53^4M^ ΔTAD can form droplet-like DPICs with random dsDNA

While wild-type (WT) human p53 assembles into a tetramer consisting of two dimers (31). Each monomer contains an amino-terminal transactivation domain (TAD, 1-62), a proline-rich domain (PRD, 63-94), a sequence-specific, highly-conserved core DBD (95-292), a short intrinsically disordered hinge domain (HD, 293-324), an oligomerization domain (OD, 325-356), and a carboxyl-terminal domain (CTD, 357-393) (32) (Fig. 1a(i)). Both the core DBD and CTD, possess DNA-binding capabilities. The CTD is responsible for nonspecific DNA searching, while the core DBD selectively binds to p53-binding motifs (33, 34). These specific dsDNA motifs are approximately 20 base pairs (bps) and contain two 10-bp half-site sequences. Each dimer within the p53 tetramer binds to one half-site sequence. The interaction of p53 with its target sequences brings about substantial alterations in downstream gene transcription, exerting a significant impact on cellular functions. For instance, binding to the p21 binding site, a crucial p53-binding motif, can induce cell cycle arrest (31, 35). The interactions between dsDNA and p53 are finely regulated, rendering p53 an exemplary candidate for unraveling the molecular mechanisms underlying morphology transition.

**Fig. 1.**
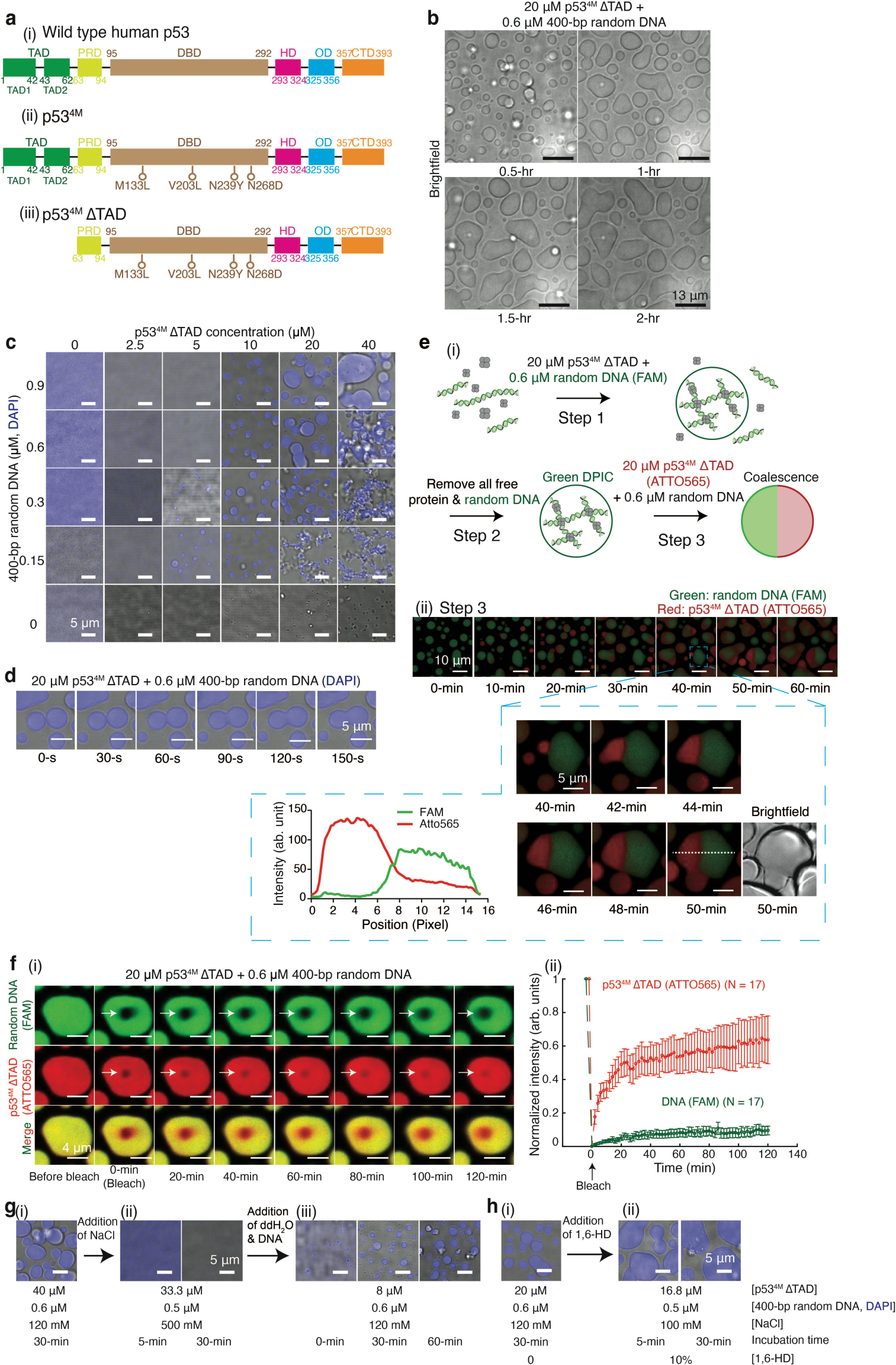
Formation of droplet-like DPICs by p53^4M^ ΔTAD with random dsDNA. (**a**) Schematic representation of WT human p53 (i), p53^4M^ (ii), and p53^4M^ ΔTAD (iii). (**b**) Time course of the fusion event between 20 μM p53^4M^ ΔTAD and 0.6 μM 400-bp random DNA (Supplementary Movie 1), with time indicated in hours. (**c**) *In vitro* droplet assays mixing 0, 0.15, 0.3, 0.6, and 0.9 μM 400-bp random DNA with 0, 2.5, 5, 10, 20, and 40 μM dark p53^4M^ ΔTAD, where DNA was labeled with DAPI. (**d**) Time course of the fusion event between two droplet-like DPICs from (b), with time indicated in seconds. (**e**) Two-color experiment. (i) Schematic of the process: Step 1, formation of green DPICs by mixing 20 μM dark p53^4M^ ΔTAD and 0.6 μM 400-bp random DNA labeled with FAM; Step 2, removal of free protein and DNA by washing; Step 3, injection of a new sample containing 20 μM p53^4M^ ΔTAD labeled with ATTO565 and 0.6 μM dark 400-bp random DNA. (ii) Time course of coalescence of two color-labeled DPICs, with time indicated in minutes. The box zooms in on a representative fusion event between 40 and 50 min, and fluorescence intensities at 50 min are plotted. (**f**) FRAP assay. (i) DPICs formed by 0.6 μM 400-bp random DNA labeled with FAM and 20 μM p53^4M^ ΔTAD labeled with ATTO565, with a 2-hour incubation and 2-hour recovery process recorded. (ii) FRAP curves with red indicating p53^4M^ ΔTAD (ATTO565) and green indicating DNA (FAM). Seventeen independent DPICs were plotted (N = 17). Error bars represent mean ± s.d. (**g**) (i) Formation of a droplet-like DPIC with 40 μM p53^4M^ ΔTAD and 0.6 μM 400-bp random DNA under 120 mM NaCl; (ii) Addition of NaCl to 500 mM and incubation for 5 min or 30 min at room temperature; (iii) Addition of water and DNA to decrease the salt concentration back to 120 mM NaCl and incubation for 5 min or 30 min at RT. (**h**) (i) Formation of a droplet-like DPIC with 20 μM p53^4M^ ΔTAD and 0.6 μM 400-bp random DNA under 0% 1,6-hexanediol (1,6-HD); (ii) Addition of 1,6-HD to 10% and incubation for 5 min or 30 min at room temperature. The working buffer for in vitro droplet experiments was 8 mM Tris-HCl (pH 7.5), 120 mM NaCl, 4% Glycerol, and 16 mM DTT. The working buffer contained no crowding agents in this study. Independent *in vitro* droplet experiments in b to h were repeated three times (n = 3) respectively.

WT human p53 exhibits susceptibility to *in vitro* degradation, the Fersht lab has identified a variant with enhanced thermodynamic stability through the substitution of four residues (M133L/V203A/N239Y/N268D) in the DNA-binding domain (DBD) – p53^4M^ (Fig. 1a(ii)) (36). Consequently, we undertook the *in vitro* purification of p53^4M^ (Supplementary Fig. 1a and **Methods**). Experimental validation of its activity was conducted through two distinct assays: electrophoretic mobility shift assays (EMSAs) (Supplementary Fig. 1b) and luciferase assays (Supplementary Fig. 1c). In *in vitro* droplet assays conducted under physiological conditions (8 mM Tris-HCl (pH 7.5) and 120 mM NaCl), co-condensate formation was not observed for 40 μM p53^4M^ in combination with 0.9 μM 400-bp DNA substrates containing non-specific sequences (random DNA) (Supplementary Fig. 1d). However, upon removal of the TAD (p53^4M^ ΔTAD, Fig. 1a(iii) and Supplementary Fig. 2a-b), 20 μM p53^4M^ ΔTAD demonstrated the capability to induce droplet-like DPIC formation when mixed with 0.6 μM 400-bp random DNA (Fig. 1b and Supplementary Movie 1). To scrutinize the intricate details of DPICs, we conducted *in vitro* droplet assays, systematically combining concentrations of 0, 0.15, 0.3, 0.6, and 0.9 μM 400-bp random DNA labeled with DAPI with 0, 2.5, 5, 10, 20, and 40 μM dark p53^4M^ ΔTAD, thereby constructing a phase diagram (Fig. 1c). Our findings indicate that neither 0.9 μM DNA nor 40 μM protein in isolation undergoes LLPS, confirming the distinctive nature of our co-condensates as DPICs. Therefore, we chose p53^4M^ ΔTAD as a biophysical model in this study to quantitatively explore the molecular mechanisms underlying morphology transition. It is imperative to note that, despite the significant biological relevance of the human p53, this work confines its focus to considering p53^4M^ ΔTAD as an ideal candidate for a thorough exploration of the quantitative biophysical mechanisms governing DPICs. It is pertinent to mention that the working buffer employed in our experiments does not contain crowding agents. Furthermore, the concentration of p53 referred to henceforth pertains to the monomeric concentration. In all instances during the *in vitro* droplet assays, samples were thoroughly mixed and incubated for a duration of 30 minutes prior to data acquisition, unless specified otherwise.

Remarkably, the droplet-like DPICs exhibited an intriguingly slow fusion kinetics (>150 seconds in Fig. 1d. To gain deeper insights into the prolonged fusion process, a two-color experiment was conducted (Fig. 1e(i)). Initially, 20 μM dark p53^4M^ ΔTAD was mixed with 0.6 μM 400-bp random DNA labeled by FAM, resulting in the formation of initial green DPICs. Subsequently, the buffer was exchanged with a fresh solution containing 20 μM p53^4M^ ΔTAD labeled by ATTO565 and 0.6 μM dark 400-bp random DNA. The ensuing 1-hour dynamic fusion process was observed in Fig. 1e(ii)-(iii). Notably, many red droplet-like DPICs underwent coalescence with green counterparts, establishing a distinct boundary during the fusion process. This two-color experiment conclusively verified the inability of the protein within red DPICs and the DNA within the green DPICs to mix well, emphasizing a pronounced material specificity of DPICs, which will be discussed in more detail in the following subsection.

We also conducted fluorescence recovery after photobleaching (FRAP) experiments (**Methods** and Supplementary Fig. 3) for 0.6 μM 400-bp random DNA labeled with FAM mixed with 20 μM p53^4M^ ΔTAD labeled with ATTO565 (Fig. 1f(i)). The FRAP analysis revealed a slow yet substantial recovery for p53^4M^ ΔTAD (depicted by the red curve in Fig. 1f(ii)). In parallel, the FAM-labeled random DNA exhibited a more limited recovery (illustrated by the green curve in Fig. 1f(ii)). These FRAP results robustly suggest that DNA serves as the structural framework, constituting the skeleton of DPIC, while the protein functions as the connecting joint within the DPIC framework.

To fully understand the biophysical nature of these droplet-like DPICs, we mixed 40 μM p53^4M^ ΔTAD with 0.6 μM 400-bp random DNA in a physiological buffer containing 120 mM NaCl (Fig. 1g(i)). Upon increasing the NaCl concentration to 500 mM, a noteworthy phenomenon unfolded—many DPICs dissipated within a brief 5-minute interval (Fig. 1g(ii)), and subsequent restoration was observed upon decreasing NaCl concentration back to 120 mM, albeit requiring a duration of 30 to 60 minutes (Fig. 1g(iii)). Furthermore, the impact of 1,6-hexanediol (1,6-HD) presence was examined, revealing that DPICs did not undergo disassembly even in the presence of 10% 1,6-HD (Fig. 1h). These findings robustly indicate that the DPICs are driven largely by electrostatic interactions between DNA and p53^4M^ ΔTAD, as opposed to hydrophobic interactions.

### The capacity of p53 to bridge different DNA duplexes induces the DPIC formation

Our subsequent inquiry delved into the microscopic architecture of the DPIC. Previous study (37) has demonstrated that the DBDs of KLF4 is proficient in forming DPICs with DNA, simultaneously exhibiting a special capability to bridge diverse DNA duplexes. This finding underscores a potential crucial correlation between the DNA-bridging capacity of KLF4 DBD and the formation of DPICs. In a parallel line of evidence, previous investigations (38) have revealed the ability of a p53 tetramer to bridge different DNA duplexes. Collectively, these observations led us to formulate a hypothesis positing that the capacity of p53 to bridge distinct DNA duplexes plays a pivotal role in inducing the formation of DPICs.

To support this idea, we conducted coarse-grained MD simulations using the 3SPN-AICG energy function (39-42) to analyze the structural and interaction features of DNA and p53 within a DPIC (**Supplementary Methods**). In this coarse-grained molecular model, each amino acid is simplified as one spherical bead centered around its C_α_ position. Similarly, each nucleotide is represented by three beads corresponding to the phosphate, sugar, and base, respectively (Fig. 2a(i) and Supplementary Fig. 4a). The structure-based potential AICG2+ (42) was applied to the core DBD and OD domains in order to maintain the structural stabilities of the folded domains and the tetrameric state of the p53^4M^ ΔTAD chains. The intrinsically disordered parts were modelled by using a statistical potential developed by Terakawa and Takada in a previous work (43), which has been demonstrated to be successful in reproducing the experimentally observed properties of the intrinsically disordered TAD domain of p53. Since p53 and DNA interact mainly through electrostatic interactions, we employed the Debye-Hückel model, which can reasonably describe the Coulombic screening effect of monovalent salt under different concentrations. All simulations were conducted with the GENESIS 1.7.1 package (44-46) at a temperature of 300 K controlled by the Langevin dynamics. More details of the coarse-grained simulations can be found in **Supplementary Methods**. Such coarse-grained modeling enables us to simulate large-scale biomolecular processes over longer timescales that are otherwise inaccessible in an atomistic model.

**Fig. 2.**
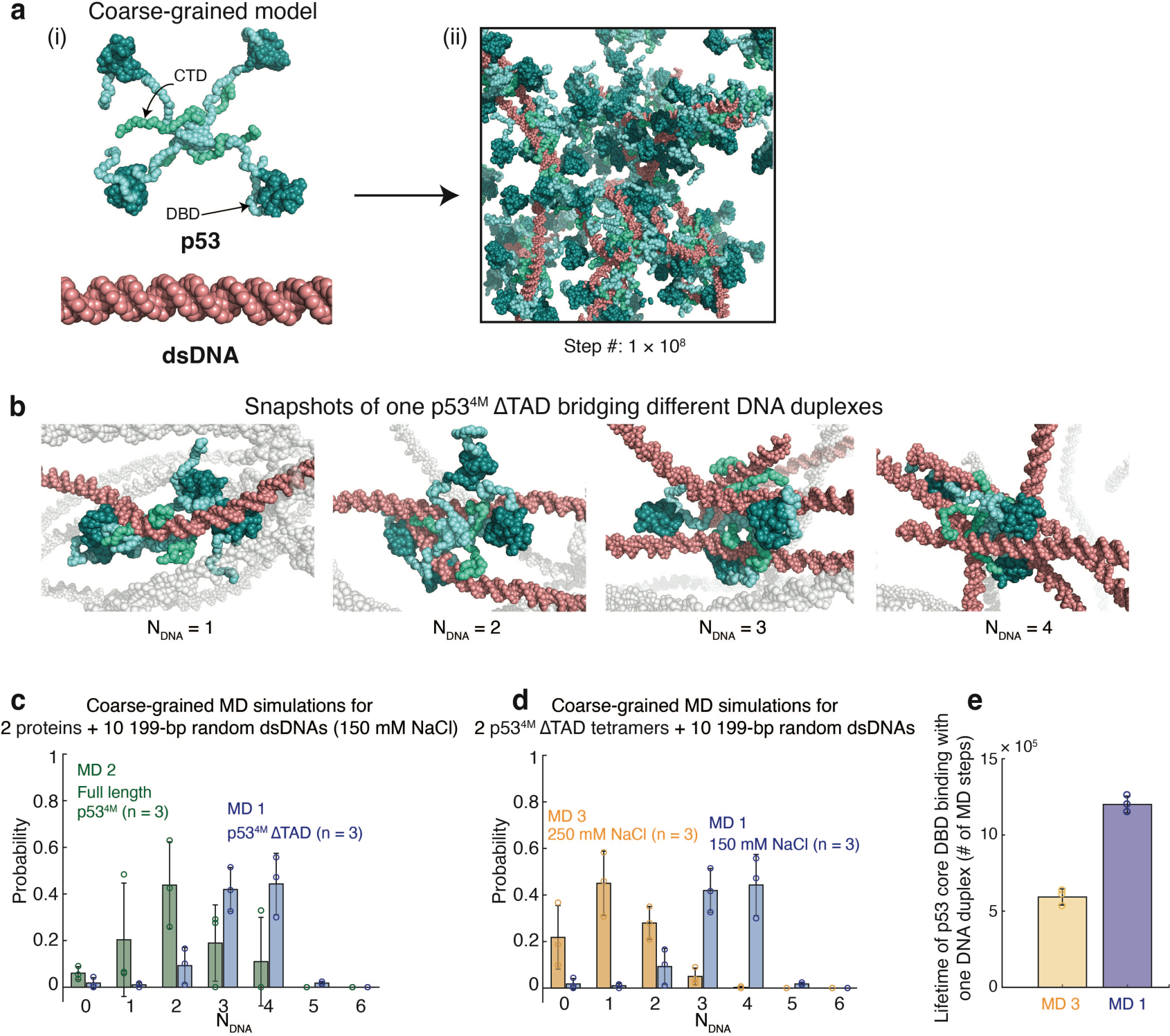
MD simulations showing the microscopic structural feature of the DPIC. (**a**) The coarse-grained MD simulations of the DPIC formation for a system containing 7 random dsDNAs and 27 p53^4M^ ΔTAD tetramers at the temperature of 300 K and the salt concentration of 150 mM. (i) The coarse-grained model of p53^4M^ ΔTAD tetramer and dsDNA chain. The DBD and CTD of p53 were colored by cyan and green, respectively. (ii) A representative structure showing the protein-DNA network formed after 1× 10^8^ steps of MD simulations. (**b**) Representative structures from the MD simulation snapshots with the p53^4M^ ΔTAD tetramer bound with one dsDNA (N_DNA_ = 1), two dsDNAs (N_DNA_ = 2), three dsDNAs (N_DNA_ = 3), and four dsDNAs (N_DNA_ = 4), respectively. The simulations were performed with the similar conditions as (a) but for a smaller system containing 2 p53^4M^ ΔTAD tetramers and 10 dsDNA chains (MD 1). (**c**) Probability distribution of the number of bound 199-bp dsDNA (N_DNA_) for each p53^4M^ ΔTAD tetramer (MD 1: blue) or full length p53^4M^ (MD 2: green) at a temperature of 300 K and a salt concentration of 150 mM. (**d**) Probability distribution of the number of bound 199-bp dsDNA for each p53^4M^ ΔTAD tetramer at a temperature of 300 K and a salt concentration of 150 mM (MD 1: blue) or 250 mM (MD 3: orange). (**e**) The average *Lifetime* of binding between p53 core DBD and DNA duplex from MD 1 and 3. The probabilities and average *Lifetime* were calculated based on the snapshots from three independent MD simulations (n = 3). The snapshots of the first 3 × 10^7^ MD steps were omitted in calculating the probability distribution of N_DNA_. Error bars, mean ± s.d.

We first conducted MD simulations for a system containing 7 random dsDNAs and 27 p53^4M^ ΔTAD tetramers (Fig. 2a(ii) and Supplementary Fig. 4b). Here the length of dsDNA was set as 199-bp in the simulations for simplicity, considering that the morphology of the DPICs does not change when the length of dsDNA increases from 199-bp to 400-bp (Supplementary Fig. 2c). This simulation began from an initial configuration in which all molecules were randomly positioned in a cubic box with rare inter-molecule bridging interactions (Supplementary Fig. 4b(i)). While the timescale of the MD simulation was still much shorter than an experimentally relevant timescale, one can observe that the DNA-p53 system gradually developed multivalent bridging interactions (Supplementary Fig. 4b(ii), c, and Supplementary Movie 2). Over time, one p53^4M^ ΔTAD tetramer has an opportunity to bind with multiple dsDNA chains, and vice versa.

The above system has a large number of degrees of freedom, and the conformational sampling in these molecular simulations is extremely slow, making quantitative analysis difficult. To more comprehensively and quantitatively characterize the effects of different factors on the capability of p53 tetramers to bridge multiple dsDNA chains, we performed extensive simulations on a much smaller system containing 2 p53^4M^ ΔTAD tetramers and 10 199-bp random dsDNA chains in a 150 mM salt concentration. This set of MD simulations was named as MD 1 (blue bars in Fig. 2c-d, and Supplementary Movie 3). In order to quantitatively characterize multivalent bridging interactions, we defined N_DNA_ to represent the number of different dsDNA chains that one p53^4M^ ΔTAD tetramer can bridge (**Supplementary Methods** and Fig. 2b). We obtained N_DNA_ = 3.5 ± 0.7 (n = 3) for MD 1 (Supplementary Fig. 5a(i)), indicating that each p53^4M^ ΔTAD tetramer is able to bridge multiple different dsDNA chains. Similar results were obtained when the number of p53^4M^ ΔTAD tetramers and dsDNA chains were altered within a reasonable range (Supplementary Fig. 5b). For comparison, we also performed two additional sets of simulations: (*i*) MD 2: 2 full-length p53^4M^ tetramers and 10 199-bp random dsDNA chains in a 150 mM salt concentration, N_DNA_ = 2.0 ± 0.9 (n = 3) (green bars in Fig. 2c, Supplementary Fig. 5a(ii), and Supplementary Movie 4); (*ii*) MD 3: 2 p53^4M^ ΔTAD tetramers and 10 199-bp random dsDNA chains in a 250 mM salt concentration, N_DNA_ = 1.1 ± 0.9 (n = 3) (orange bars in Fig. 2d, Supplementary Fig. 5a(iii), and Supplementary Movie 5). While the effective concentration and stoichiometry used in MD simulations is different from what is employed in the experimental measurements in order to improve sampling efficiency, the dramatic decrease of the protein-DNA bridging capacity due to the TAD addition (MD 2) and the salt concentration increasement (MD 3) is consistent with the above experimental observations, highlighting the critical role of p53 bridging across different DNA duplexes for DPIC formation. Remarkably, in addition to the KLF4 complexes with DNA (37), these microscopic configurations align with those observed in cohesion complexes with DNA (47), and human cGAS complexes with DNA (48).

From the above MD simulations, we also determined the average dwelling time for a p53 core DBD binding with one DNA duplex, which was named as “Lifetime of p53 core DBD binding with one DNA duplex” (Supplementary Fig. 4a and **Supplementary Methods**). The average Lifetime in MD 1 (150 mM NaCl) is (1.20 ± 0.06) × 10^6^ (n = 3) MD steps (Fig. 2e). Upon salt concentration increases to 250 mM, the value dramatically decreased to (0.59 ± 0.05) × 10^6^ (n = 3) MD steps (MD 3 in Fig. 2e), demonstrating the salt-resulted weakening of p53-dsDNA bridging interactions.

In this context, we introduce the term “ε_app_” to denote the apparent inter-molecule interaction strengths within the DPIC. Given the distinctive microscopic structure of DPIC, the number of bridges, symbolized by p53 molecules on each DNA, and the binding affinity between DNA and protein serve as two important factors governing ε_app_.

### Increasing the ε_app_ induces a morphological transition from a droplet-like to a “pearl chain”-like structure

Analysis of the phase diagram presented in Fig. 1c reveals two distinct pathways that lead to alterations in the morphology of DPICs: (i) at 20 μM p53^4M^ ΔTAD, 0.9 μM 400-bp random DNA substrates exhibit detectable droplet-like DPIC formation, whereas concentrations below 0.3 μM of DNA initiate a formation of distinguishable “pearl chain”-like DPICs; (ii) at 0.15 μM 400-bp random DNA substrates, 5 μM p53^4M^ ΔTAD shows detectable droplet-like DPIC formation, whereas concentrations exceeding 10 μM protein also initiate the formation of distinguishable “pearl chain”-like DPICs. These results suggest that altering the initial molar ratio between p53 and DNA (m = [p53] / [DNA]) can influence DPIC morphology.

This observation arises from the unique microscopic structure of DPICs. Coarse-grained MD simulations depicted in Fig. 2 suggest that the p53 capacity to bridge different DNA duplexes induces DPIC formation. Therefore, the number of bridges—represented by p53 molecules—on each DNA serves as an important factor regulating the inter-molecule interaction strengths within the DPIC system. Consequently, changing m alters the number of bridges on each DNA, resulting in a shift in DPIC morphology.

To examine this aspect carefully, we conducted two-color *in vitro* droplet assays by combining 20 μM p53^4M^ ΔTAD labeled with ATTO565 and varying concentrations of 400-bp random DNA labeled with FAM (0, 0.075, 0.15, 0.225, 0.3, 0.375, 0.45, 0.6, 0.75, 0.9, 1.2, 1.5, and 1.8 μM) (Fig. 3a). Notably, at 1.8 μM 400-bp random DNA, droplet-like DPICs were readily observable, corresponding to an m value of 20/1.8 = 11.1. However, as the DNA concentration decreased to 0.225 μM, a transition in DPIC morphology from droplet-like to “pearl chain”-like occurred, coinciding with an increase in m from 11.1 to 88.9. These outcomes strongly imply that an increase in m results in a higher number of bridges on each DNA, thereby transitioning the DPIC morphology from droplet-like to “pearl chain”-like. Furthermore, there exists a threshold for m (m_th_), approximately 66.7, beyond which “pearl chain”-like DPIC formation is induced.

**Fig. 3.**
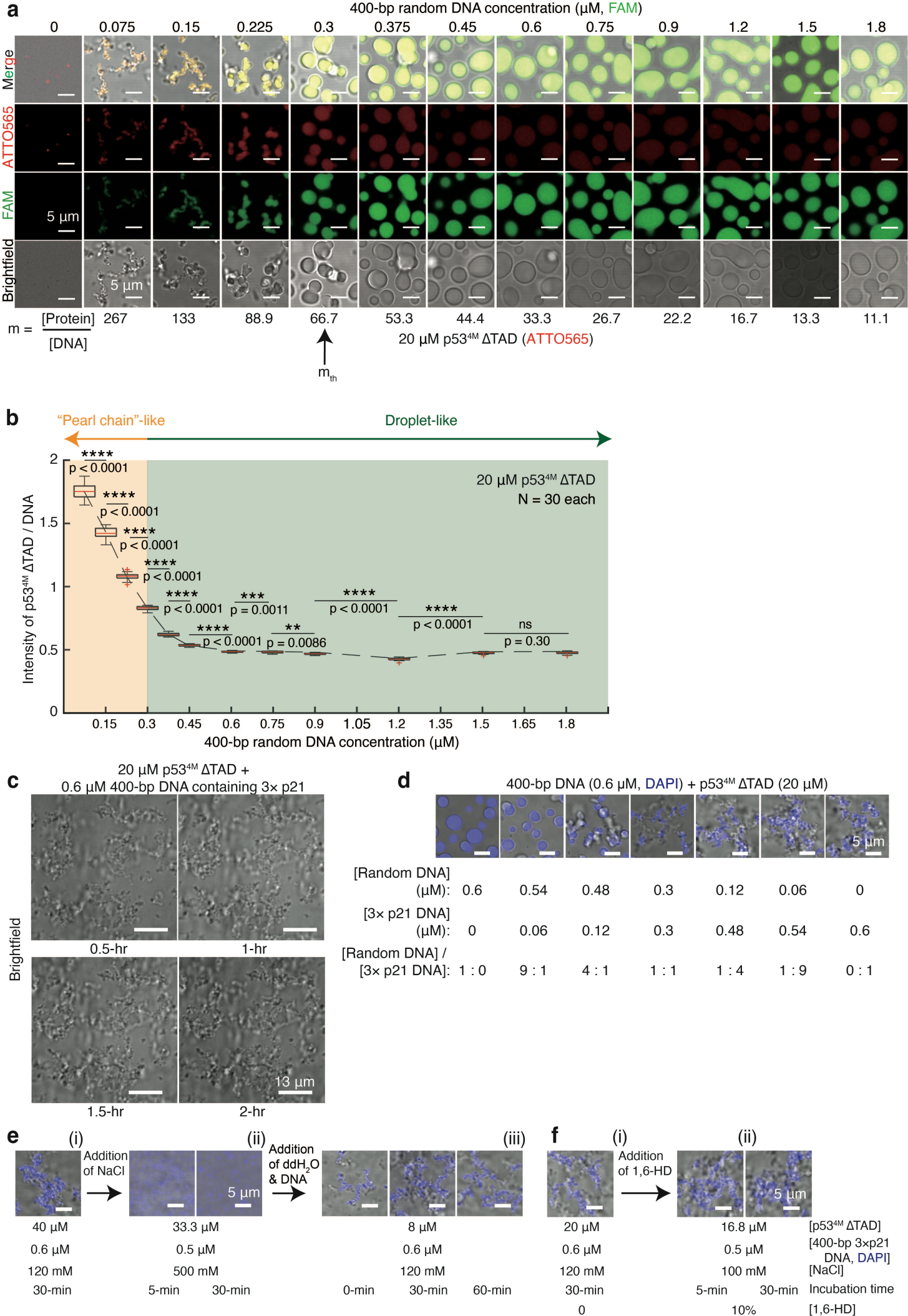
Morphological transition of DPICs from droplet-like to "pearl chain"-like induced by increasing the total interaction strengths of the system. (**a**) Two-color *in vitro* droplet assays involving the mixing of 0, 0.075, 0.15, 0.225, 0.3, 0.375, 0.45, 0.6, 0.75, 0.9, 1.2, 1.5, and 1.8 μM 400-bp random DNA labeled by FAM with 20 μM p53^4M^ ΔTAD labeled by ATTO565. The initial molar ratio between p53 and DNA for each condition (m value) is provided. Independent *in vitro* droplet experiments were conducted three times (n = 3). (**b**) Intensity ratio of p53^4M^ ΔTAD / DNA within DPIC for all conditions in (a). The total number (N) of DPICs examined for each condition was 30. For the boxplot, the red bar represents median. The bottom edge of the box represents 25^th^ percentiles, and the top is 75^th^ percentiles. Most extreme data points are covered by the whiskers except outliers. The ‘+’ symbol is used to represent the outliers. Statistical significance was analyzed using unpaired t test for two groups. P value: two-tailed; p value style: GP: 0.1234 (ns), 0.0332 (*), 0.0021 (**), 0.0002 (***), <0.0001 (****). Confidence level: 95%. (**c**) Four panels depicting the time course of fusion events in an *in vitro* droplet experiment with 20 μM p53^4M^ ΔTAD and 0.6 μM 400-bp DNA containing 3× p21 binding motifs (400-bp 3× p21 DNA) (Supplementary Movie 6), with time indicated at the bottom in hours. (**d**) *In vitro* droplet experiments by mixing 400-bp random DNA and 400-bp DNA containing 3× p21 binding motifs with 20 μM dark p53^4M^ ΔTAD, with changing molar ratios of 400-bp random DNA over 400-bp 3× p21 DNA. (**e**) (i) “Pearl chain”-like DPIC formation with 40 μM p53^4M^ ΔTAD and 0.6 μM 400-bp DNA containing 3× p21 binding motifs under 120 mM NaCl conditions. (ii) Addition of NaCl to 500 mM and incubation for 5 min or 30 min at room temperature (RT). (iii) Addition of extra water and DNA to reduce salt concentration back to 120 mM NaCl and incubation for 0 min, 30 min, or 60 min at RT. (**f**) (i) “Pearl chain”-like DPIC formation with 20 μM p53^4M^ ΔTAD and 0.6 μM 400-bp DNA containing 3× p21 binding motifs under 1,6-HD conditions of 0% 1,6-HD. (ii) Addition of 1,6-HD to 10% and incubation for 5 min or 30 min at RT. DNA was labeled with DAPI. Independent *in vitro* droplet experiments in (a and c-f) were repeated three times (n = 3) each.

To assess the actual molar ratio within the DPIC, we measured the intensity ratio (Intensity of p53^4M^ ΔTAD (ATTO565) / DNA (FAM)). Our findings revealed that the ratio for droplet-like DPICs is substantially lower than that of “pearl chain”-like DPICs (Fig. 3b), confirming that the number of bridges on each DNA is an important factor governing intermolecular interaction strengths within the DPIC system.

EMSAs revealed a significantly enhanced binding affinity of p53^4M^ ΔTAD to the p21 binding motifs in comparison to random DNA (Supplementary Fig. 2b). Subsequently, we explored whether the addition of p21 binding motifs into the DNA sequence could also modulate the morphology of DPICs. We devised new *in vitro* droplet assays. In contrast to the droplet-like morphology observed with 400-bp random DNA in conjunction with p53^4M^ ΔTAD (Fig. 1b(i) and Supplementary Movie 1), the combination of 0.6 μM 400-bp DNA containing 3× p21 binding motifs with 20 μM p53^4M^ ΔTAD also facilitated the formation of “pearl chain”-like DPICs (Fig. 3c and Supplementary Movie 6). A comprehensive examination of these “pearl chain”-like co-condensates was undertaken through: (*i*) *in vitro* droplet assays involving the mixing of 0, 0.15, 0.3, 0.6, and 0.9 μM 400-bp DNA containing 3× p21 binding motifs with 10, 20, and 40 μM dark p53^4M^ ΔTAD (Supplementary Fig. 6a); (*ii*) *in vitro* droplet assays incorporating the mixing of 0.6 μM 400-bp DNA containing varying numbers of p21 binding motifs (0×, 1×, 2×, 3×, and 4×) with 0, 10, 20, and 40 μM dark p53^4M^ ΔTAD (Supplementary Fig. 6b). The results demonstrated a “pearl chain”-like morphology in DPICs when the DNA substrate carried multiple copies of the p21 sequence. To observe the morphological transition, we mixed 400-bp DNA containing 3× p21 binding motifs with 400-bp random DNA, generating 0.6 μM 400-bp DNA substrates with varying concentrations of p21 sequences (Fig. 3d). At 20 μM p53^4M^ ΔTAD, 0.6 μM DNA substrates devoid of p21 exhibited detectable droplet-like DPIC formation. Beyond a concentration of 0.3 μM for 400-bp DNA containing 3× p21 binding motifs, these DNA substrates initiated the formation of discernible “pearl chain”-like DPICs. These findings confirm that increasing the binding affinity between dsDNA and protein can also induce a morphological shift from droplet-like to “pearl chain”-like.

Analogous to the droplet-like DPICs induced by random DNA, the “pearl chain”-like DPIC is profoundly responsive to NaCl concentration but remains unaffected by the presence of 1,6-HD (Fig. 3e-f). This outcome underscores that the formation of “pearl chain”-like DPICs is also predominantly driven by electrostatic interactions rather than hydrophobic interactions.

Taken together, varying the initial molar ratio between p53 and DNA, as well as manipulating the DNA sequence, emerge as two experimental approaches for fine-tuning the ε_app_. Increasing the ε_app_ induces a morphological transition from a droplet-like to a “pearl chain”-like structure.

In addition to the incorporation of p21 binding motifs, manipulating salt concentration, which can regulate the binding affinity between DNA and protein, can also alter the ε_app_. Notably, a reduction in salt concentration from 250 mM to 120 mM induced a morphological transition in DPICs formed by 0.15 μM 400-bp random DNA mixed with 20 μM p53^4M^ ΔTAD, shifting from a droplet-like to a “pearl chain”-like configuration (Supplementary Fig. 2d). In conclusion (Fig. 1 and 3), our results emphasize that increasing the ε_app_ induces a morphological transition from a droplet-like to a “pearl chain”-like structure

### Droplet-like DPICs exhibit different types of viscoelastic properties

Recent literature extensively discusses the viscoelastic nature of nucleic acid and protein co-condensation, both *in vitro* (14, 15) and *in vivo* (49). Our fusion experiments and FRAP experiments (Fig. 1d-f) consistently point towards distinctive material characteristics of DPICs. Leveraging the atomic force microscopy-based force spectroscopy (AFM-FS) technique, which has proven effective in probing the mechanical properties of biomolecular condensates, we conducted measurements to unveil the material properties of DPIC (**Methods** and Fig. 4a(i)).

**Fig. 4.**
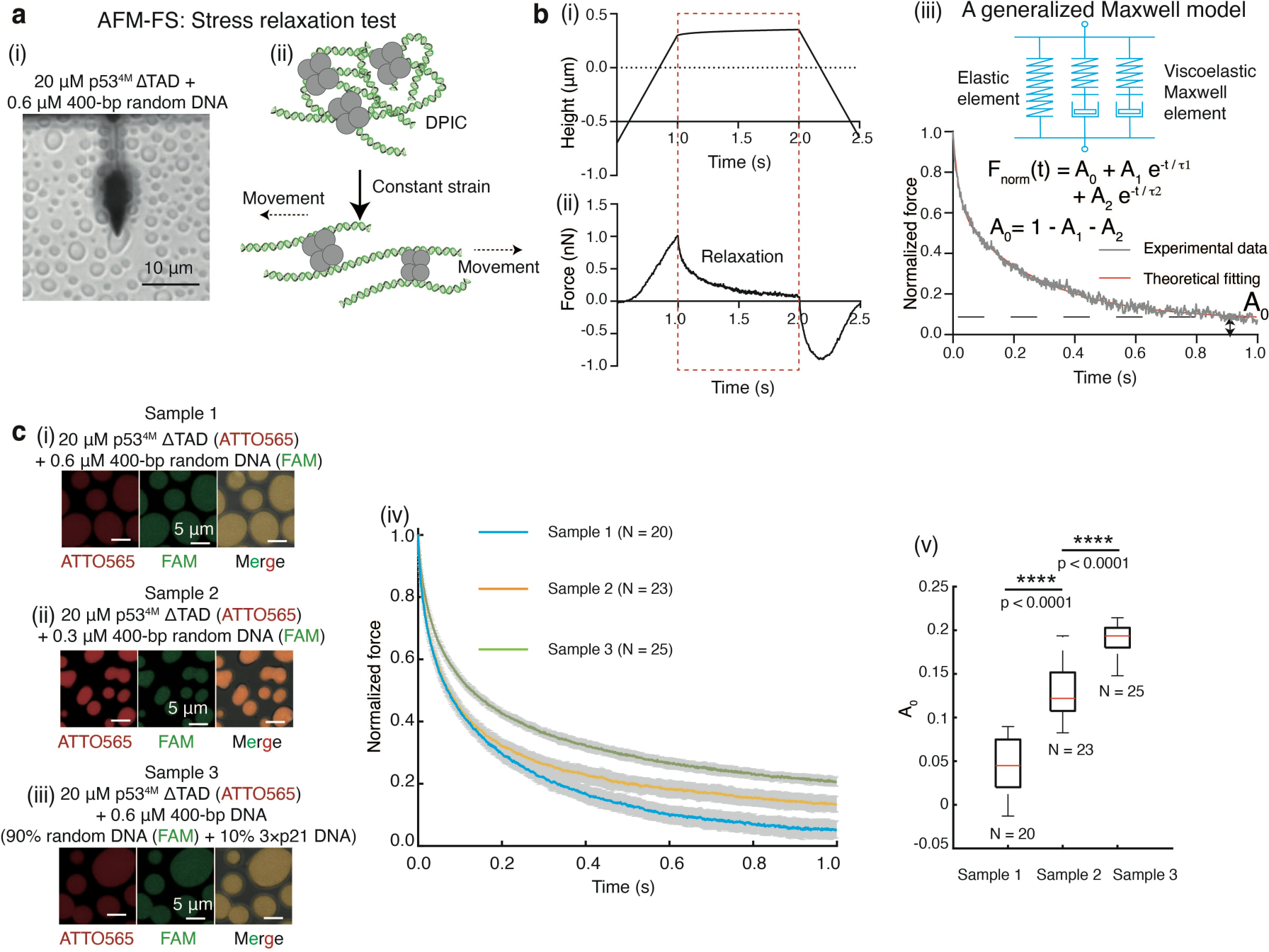
Characterizing the viscoelastic behavior of DPICs using AFM-FS. (**a**) AFM-FS experiments conducted on the droplet-like DPIC composed of 20 μM p53^4M^ ΔTAD and 0.6 μM 400-bp random DNA. (i) Top-view image of the DPIC under an AFM probe. (ii) Schematic illustration of the stress relaxation test. (**b**) Representative stress relaxation curve of the DPIC. (i) Height-time curve. (ii) Force-time curve. The orange dotted box indicates the relaxation process, where the probe’s height remained constant, and the force decreased. (iii) Schematic diagrams of a generalized Maxwell model, comprising one spring and two Maxwell elements in parallel, used for fitting the normalized relaxation curves. (**c**) *In vitro* droplet experiments by combining 20 μM p53^4M^ ΔTAD and 0.6 μM 400-bp random DNA (Sample 1) (i), 20 μM p53^4M^ ΔTAD and 0.3 μM 400-bp random DNA (Sample 2) (ii), and 20 μM p53^4M^ ΔTAD and 400-bp DNA containing 90% 400-bp random DNA and 10% 400-bp DNA containing 3× p21 binding motif (Sample 3) (iii). p53^4M^ ΔTAD was labeled with ATTO565, and 400-bp DNA was labeled with FAM. DPICs were incubated for 1 hour at room temperature. (iv) Normalized relaxation curves of the DPICs: Sample 1, blue; Sample 2, orange; Sample 3, green. The colored line represents the mean value, and the gray area represents s.d. (v) Boxplot of A_0_ for the three samples in c(i). The total number (N) of DPICs examined in a single AFM-FS experiment: N = 20 for Sample 1; N = 23 for Sample 2; N = 25 for Sample 3. For the boxplot, the red bar represents median. The bottom edge of the box represents 25^th^ percentiles, and the top is 75^th^ percentiles. Most extreme data points are covered by the whiskers except outliers. The ‘+’ symbol is used to represent the outliers. Statistical significance was analyzed using unpaired t test for two groups. P value: two-tailed; p value style: GP: 0.1234 (ns), 0.0332 (*), 0.0021 (**), 0.0002 (***), <0.0001 (****). Confidence level: 95%.

To quantitatively assess viscoelasticity, we conducted an AFM-FS experiment known as the stress relaxation test (Fig. 4a(ii)) (50). In these experiments, the AFM probe descended to a specific height (∼0.3 μm in Fig. 4b(i)) for 1 second, capturing a relaxation curve of force over this period (Fig. 4b(ii)). We employed a generalized Maxwell model featuring one spring and two Maxwell elements in parallel to fit these normalized relaxation curves (**Supplementary Methods**, Fig. 4b(iii), and Supplementary Fig. 7) (51, 52). The spring contributes a pure elastic effect, evident in the A_0_ term. The two Maxwell elements represent two relaxation stages, where A_1_ and A_2_ correspond to the force decay of the elements, and τ_1_ and τ_2_ represent the relaxation time. The relaxation time correlates with the viscosity of the DPIC. Specifically, A_0_ = 1 – A_1_ – A_2_. When A_0_ = 0, the DPIC behaves as a viscoelastic fluid. Conversely, when A_0_ > 0, the DPIC exhibits viscoelastic solid characteristics (50). Due to methodological constraints, AFM-FS experiments could only be conducted on samples exhibiting a droplet-like DPIC morphology (Fig. 4c).

We initiated the stress relaxation test for droplet-like DPICs composed of 0.6 μM 400-bp random DNA mixed with 20 μM p53^4M^ ΔTAD (Fig. 4c(i)), and the stress relaxation curve was subsequently normalized (the blue curve in Fig. 4c(iv)). In this set of experiments, a total of 20 independent DPICs were measured. The values obtained are as follows: A_0_ of 0.04 ± 0.03 (mean ± s.d., N = 20), τ_1_ of 0.26 ± 0.03 seconds (mean ± s.d., N = 20), and τ_2_ of 0.022 ± 0.003 seconds (mean ± s.d., N = 20). Given that A_0_ is close to zero, this outcome implies that this particular DPIC exhibits viscoelastic fluid-like behavior.

Using this condition as a reference, we conducted two additional AFM-FS experiments: (*i*) examining droplet-like DPICs comprising 0.3 μM 400-bp random DNA mixed with 20 μM p53^4M^ ΔTAD (Fig. 4c(ii)), with the generalized Maxwell model (Fig. 4b(iii)) applied to fit the curve (orange curve in Fig. 4c(iv)): A_0_ of 0.13 ± 0.03 (mean ± s.d., N = 23), τ_1_ of 0.28 ± 0.08 seconds (mean ± s.d., N = 23), and τ_2_ of 0.028 ± 0.005 seconds (mean ± s.d., N = 23); (*ii*) investigating droplet-like DPICs composed of 0.6 μM 400-bp DNA (90% random DNA and 10% DNA containing 3× p21 binding motifs) mixed with 20 μM p53^4M^ ΔTAD (Fig. 4c(iii)), with the generalized Maxwell model (Fig. 4b(iii)) employed to fit the curve (green curve in Fig. 4c(iv)): A_0_ of 0.19 ± 0.02 (mean ± s.d., N = 25), τ_1_ of 0.33 ± 0.04 seconds (mean ± s.d., N = 25), and τ_2_ of 0.030 ± 0.004 seconds (mean ± s.d., N = 25). These findings indicate that A_0_ is greater than zero in both conditions (Fig. 4c(v)), corresponding a viscoelastic solid regime.

Thus, droplet-like DPICs can exhibit diverse viscoelastic properties, demonstrating features characteristic of both viscoelastic fluids and viscoelastic solids. Furthermore, these findings suggest that increasing the number of bridges on each DNA molecule or enhancing the binding affinity between DNA and protein—thus increasing the ε_app_ within DPICs—may trigger a phase transition from a viscoelastic fluid to a viscoelastic solid state in droplet-like DPICs.

### Two types of transitions in the two-component phase diagram of DPICs combining p53^4M^ ΔTAD and 400-bp random DNA

To delineate the molecular mechanism governing condensate morphology, we initially inquire into the relationship between material property transition, morphological transition, and phase separation. Addressing this question requires the construction of a phase diagram for two-component DNA-protein cophase separation.

We conducted three sets of two-color *in vitro* droplet assays: (i) Combining 20 μM p53^4M^ ΔTAD labeled with ATTO565 with varying concentrations of 400-bp random DNA labeled with FAM (ranging from 0, 0.075, 0.15, 0.225, 0.3, 0.375, 0.45, 0.6, 0.75, 0.9, 1.2, 1.5, to 1.8 μM) (Fig. 3a); (ii) Combining 5 μM p53^4M^ ΔTAD labeled with ATTO565 with varying concentrations of 400-bp random DNA labeled with FAM (ranging from 0, 0.075, 0.15, 0.225, 0.3, 0.45, 0.6, to 0.9 μM) (Fig. 5a); (iii) Combining 0.075 μM 400-bp random DNA labeled with FAM with varying concentrations of p53^4M^ ΔTAD labeled with ATTO565 (ranging from 0, 2.5, 5, to 10 μM) (Fig. 5b). By measuring the intensity of p53^4M^ ΔTAD (ATTO565) and 400-bp random DNA (FAM) in both the dense and dilute phases, we generated a two-component phase diagram of DPIC (Fig. 5c). The coexistence region is centrally located, encompassing all tie lines connecting both the dense and dilute phases for each condition. Interestingly, all tie lines exhibit positive slopes, indicative of cooperativity between the components (53).

**Fig. 5.**
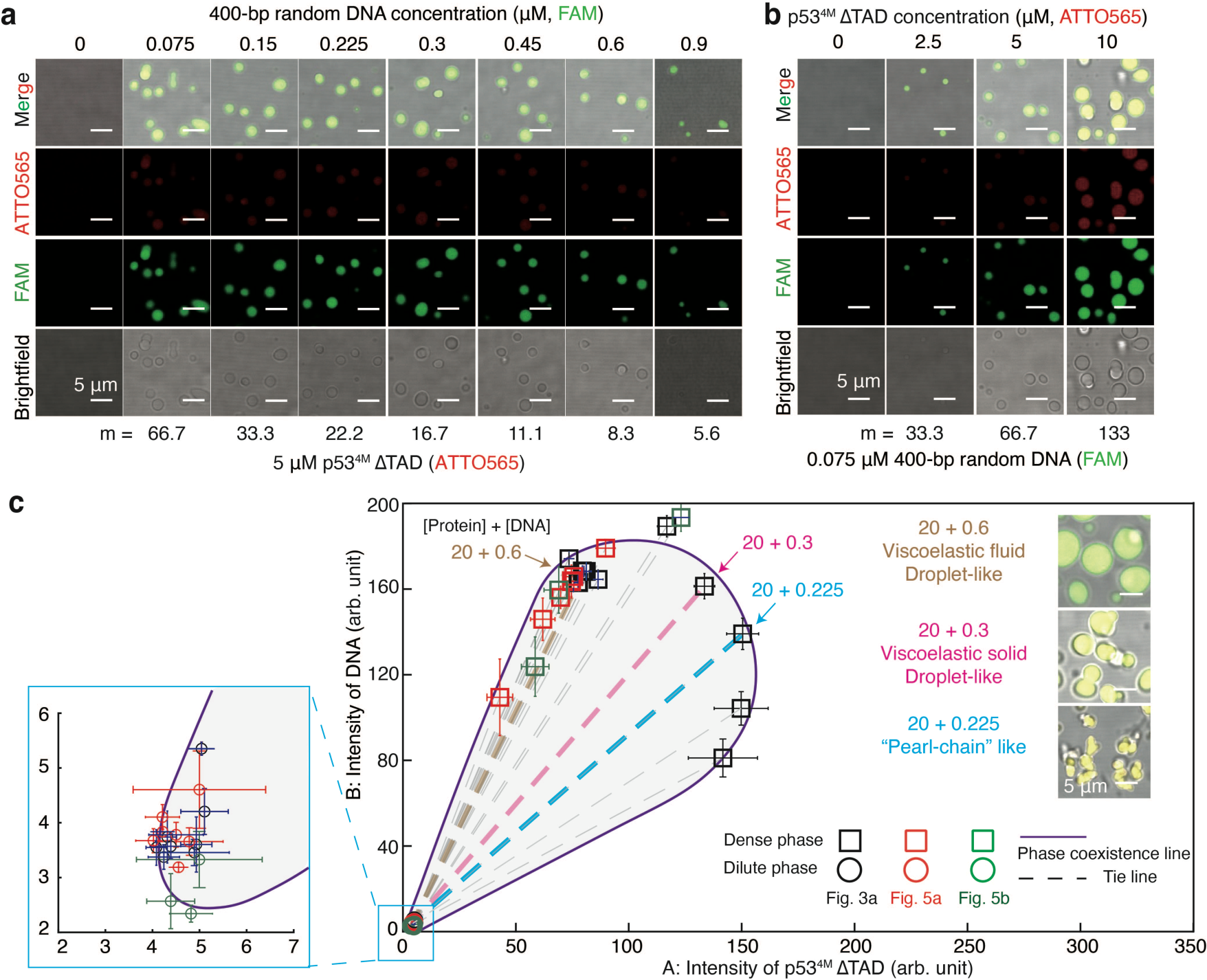
Two types of transitions in the two-component phase diagram of DPICs. (**a**) Two-color *in vitro* droplet assays were conducted by combining 0, 0.075, 0.15, 0.225, 0.3, 0.45, 0.6, and 0.9 μM 400-bp random DNA labeled with FAM with 5 μM p53^4M^ ΔTAD labeled with ATTO565. (**b**) Two-color *in vitro* droplet assays involved mixing 0, 2.5, 5, and 10 μM p53^4M^ ΔTAD labeled with ATTO565 with 0.075 μM 400-bp random DNA labeled with FAM. The working buffer for *in vitro* droplet assays comprised 8 mM Tris-HCl (pH 7.5), 120 mM NaCl, 4% Glycerol, and 16 mM DTT. Independent *in vitro* droplet experiments were performed three times (n = 3). (**c)** Two-component phase diagram of DPIC. The intensity of p53^4M^ ΔTAD (ATTO565) and 400-bp random DNA (FAM) in both the dense (square) and dilute phases (circle) from all conditions in Fig. 3a (black), 5a (red), and 5b (green) were plotted. The box zooms in on all data points in the dilute phase. The numbers labeling the dense or dilute phase represent the initial concentration of p53^4M^ ΔTAD and the initial concentration of 400-bp random DNA. The dashed line connecting data points of the dense phase and dilute phase for each condition represents the tie line.

The slopes of these tie lines exhibit interesting variation (Supplementary Fig. 8a-c), delineating two distinct thresholds indicative of different transitions. Our AFM-FS experiments revealed that despite both types of DPICs, one composed of 0.6 μM 400-bp random DNA mixed with 20 μM p53^4M^ ΔTAD (Fig. 4c(i)) and the other of 0.3 μM 400-bp random DNA mixed with 20 μM p53^4M^ ΔTAD (Fig. 4c(ii)), exhibiting a droplet-like morphology, the former, with 0.6 μM DNA, displays characteristics of a viscoelastic fluid, while the latter, with 0.3 μM DNA, exhibits properties of a viscoelastic solid. Thus, the first slope threshold, around 2.2, which is the slope of DPIC formed by 0.6 μM 400-bp random DNA mixed with 20 μM p53^4M^ ΔTAD (orange line in Supplementary Fig. 8d), signifies a phase transition between viscoelastic fluid and viscoelastic solid. The second slope threshold, which is located at the DPIC consisting of 0.225 μM 400-bp random DNA mixed with 20 μM p53^4M^ ΔTAD, approximately 0.9 (blue line in Supplementary Fig. 8d and Fig. 3a), indicates a morphology transition between DPICs with droplet-like morphology and those with “pearl chain”-like morphology.

### Droplet-like and “pearl chain”-like DPICs share the same size of the initial CMC stage at nanometer scale

To unravel the molecular mechanism dictating condensate morphology, we initially addressed the phase diagram of DPICs, as previously discussed. Our inquiry now shifts towards the second key question. Previous investigations (10, 14, 30) have demonstrated that increasing the ε_app_ correlates with prolonged relaxation times, thereby influencing the morphology and material properties of NAPCs. However, the question of how to connect the microscopic parameter ε_app_ with macroscopic morphology and material properties remains unanswered. Addressing this inquiry requires elucidating how intermolecular interaction strengths at the nanometer scale impact the morphological transition of condensates at the micrometer scale. Hence, in this session, we have designed new type of experiments to explore the growth dynamics of DPICs.

In previous experiments, we found that DPIC is mainly formed by fusion (Fig. 1b and e). So, we define the basic unit of DPIC fusion as critical microscopic cluster (CMC). Drawing from the theory of phase separation (54), we posited an assumption: when the initial concentrations of protein and DNA decrease to saturation concentration (*C_sat_*), the total volume fraction of DPICs becomes exceedingly small, and the size of a single DPIC corresponds to a CMC. In this section, we want to explore whether we could quantitatively measure and compare the size of the CMCs for both droplet-like and “pearl-chain”-like DPICs.

The spatial constraints imposed by the optical diffraction limit (∼200 nm) in confocal fluorescence microscopy for *in vitro* droplet assays present a challenge in measuring nanometer-sized CMCs (Supplementary Fig. 2e). To accurately quantify the CMCs, we employed a confocal-based fluorescence fluctuation microscopy – the dual-color fluorescence cross-correlation spectroscopy (dcFCCS) assay (Fig. 6a) (55, 56). We labeled p53^4M^ ΔTAD and 400-bp DNAs with Cyanine5 (Cy5) and FAM, respectively (Fig. 6a(i)). These labeled components freely diffused through the confocal detecting volume (Fig. 6a(ii)). Upon protein-DNA heterocomplex formation, the signals from protein and DNA contributed to the cross-correlation curves, and the relaxation times of these curves facilitated the precise quantification of the hydrodynamic radii of DPICs (Fig. 6a(iii)).

**Fig. 6.**
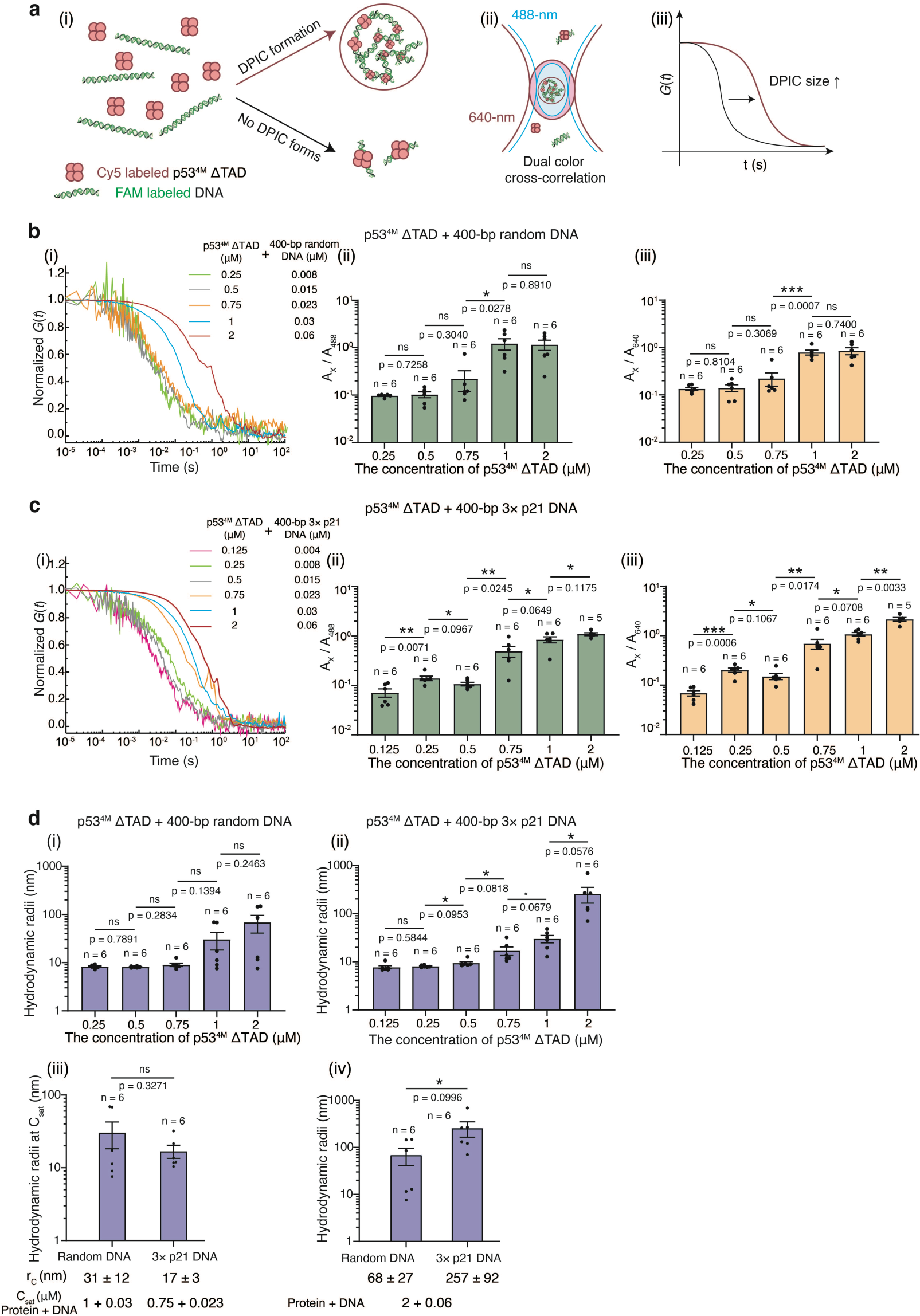
Droplet-like and “pearl chain”-like DPICs can form similar nm-sized CMCs. (**a**) Schematic diagram of the dcFCCS assay. (i)-(ii) p53^4M^ ΔTAD labeled with Cy5 and 400-bp DNA labeled with FAM (i) diffuse freely through the detection volumes of 488- and 640-nm lasers (ii). (iii) Only the heterocomplexes carrying both Cy5-labeled protein and FAM-labeled DNA contribute to cross-correlation curves, with relaxation times correlating to their size. (**b-c**) dcFCCS analysis of conditions involving the mixing of p53^4M^ ΔTAD and 400-bp DNA. (i) Representative normalized dcFCCS curves for mixtures of Cy5-labeled p53^4M^ ΔTAD and FAM-labeled 400-bp DNA, with listed concentrations of protein and DNA. (ii) and (iii) A_X_ / A_488_ and A_X_ / A_640_ indicate the participation ratios of p53^4M^ ΔTAD and 400-bp DNA, respectively, in the protein-DNA heterocomplexes. 400-bp random DNA (b) and 400-bp DNA containing 3× p21 binding motifs (3× p21 DNA) (c). (**d**) Hydrodynamic radii of heterocomplexes formed by p53^4M^ ΔTAD and (i) 400-bp random DNA or (ii) 400-bp 3× p21 DNA. (iii) The hydrodynamic radii of heterocomplexes containing these two types of DNA substrates under C_sat_ conditions, estimating the size of CMC of DPICs (r_c_). (iv) The hydrodynamic radii of heterocomplexes containing these two types of DNA substrates under the condition composed of 2 μM p53^4M^ ΔTAD and 0.06 μM DNA. The working buffer for dcFCCS assays was 8 mM Tris-HCl (pH 7.5), 120 mM NaCl, and 16 mM DTT. Independent dcFCCS assays were performed more than 5 times for each condition. Error bars in Fig. 5b, c, d, represent mean ± s.e.m. Statistical significance was analyzed using unpaired t test for two groups. P value: two-tailed; p value style: GP: 0.1234 (ns), 0.0332 (*), 0.0021 (**), 0.0002 (***), <0.0001 (****). Confidence level: 95%.

Maintaining a fixed ratio of p53^4M^ ΔTAD to 400-bp DNA concentrations at 20:0.6, we conducted a series of experiments with increasing concentration (Fig. 6b(i)). For random DNA, known to induce droplet-like DPICs (Fig. 1b), when the concentrations of p53^4M^ ΔTAD and DNA were below 1 μM and 0.03 μM, respectively, the participation ratios of p53^4M^ ΔTAD and DNA were notably low (Fig. 6b(ii)-(iii)). This suggests that most proteins and DNAs did not partake in protein-DNA heterocomplex formation. However, with concentrations of p53^4M^ ΔTAD and DNA at 1 μM and 0.03 μM or higher, respectively, both the participation ratios and hydrodynamic radii of heterocomplex showed a significant increase (Fig. 6b(ii)-(iii) and d(i)). The sharp rise in the participation ratio at 1 μM p53^4M^ ΔTAD and 0.03 μM 400-bp random DNA indicates this as the *C_sat_* condition for droplet-like DPICs.

Similarly, for DNA containing 3× p21 binding motifs, capable of inducing “pearl chain”-like DPICs (Fig. 3c), increasing protein and DNA concentrations resulted in elevated participation ratios and hydrodynamic radii of protein-DNA heterocomplex (Fig. 6c and d(ii)). The participation ratios of proteins and DNAs significantly increased at 0.75 μM p53^4M^ ΔTAD and 0.023 μM 400-bp DNA containing 3× p21 binding motifs, marking this as the *C_sat_* condition for “pearl chain”-like DPIC. At their respective *C_sat_* conditions, there was no significant difference in the hydrodynamic radius between droplet-like DPICs and “pearl chain”-like DPICs (Fig. 6d(iii)). This result suggests that both droplet-like DPICs and “pearl chain”-like DPICs encompass a stage of CMCs with a consistent size of about 15-30 nm. Additionally, the noted CMC size of DPICs containing cGAS and dsDNA is approximately 70 nm (56), indicating potential variability in CMC sizes across different types of DPICs.

Remarkably, even slight increases in the concentration of protein and DNA beyond the saturation (*C_sat_*) conditions, such as 0.06 μM 400-bp DNA and 2 μM p53^4M^ ΔTAD, yielded a fourfold difference in hydrodynamic radius between droplet-like and "pearl chain"-like DPICs (Fig. 6d(iv)). This observation strongly suggests that although droplet-like and “pearl chain”-like DPICs possess the same nanometer-scale CMCs under saturation conditions (Fig. 6d(iii)), the manner in which these CMCs aggregate into larger droplet-like or “pearl chain”-like DPICs under higher concentration conditions may diverge significantly.

### The growth dynamics of DPICs

Integrating both AFM-FS and dcFCCS experiments (Fig. 4 and 6), we have developed a model to elucidate the growth dynamics of DPICs (Fig. 7b). Our model delineates two distinct stages in the DPIC formation process. In the initial stage (Stage I), interactions between DNA and protein give rise to CMCs with a radius of approximately 15-30 nm. These CMCs, exhibiting varying ε_app_, may manifest diverse relaxation times and viscoelastic properties (Fig. 6 and 7b(i)). Subsequently (Stage II), these CMCs coalesce into larger DPICs. CMCs with more fluid-like viscoelastic (or less viscoelastic solid-like) properties readily fuse to form droplet-like DPICs, while those with more solid-like viscoelastic properties resist fusion, resulting in the formation of “pearl chain”-like DPICs (Fig. 7b(ii)).

**Fig. 7.**
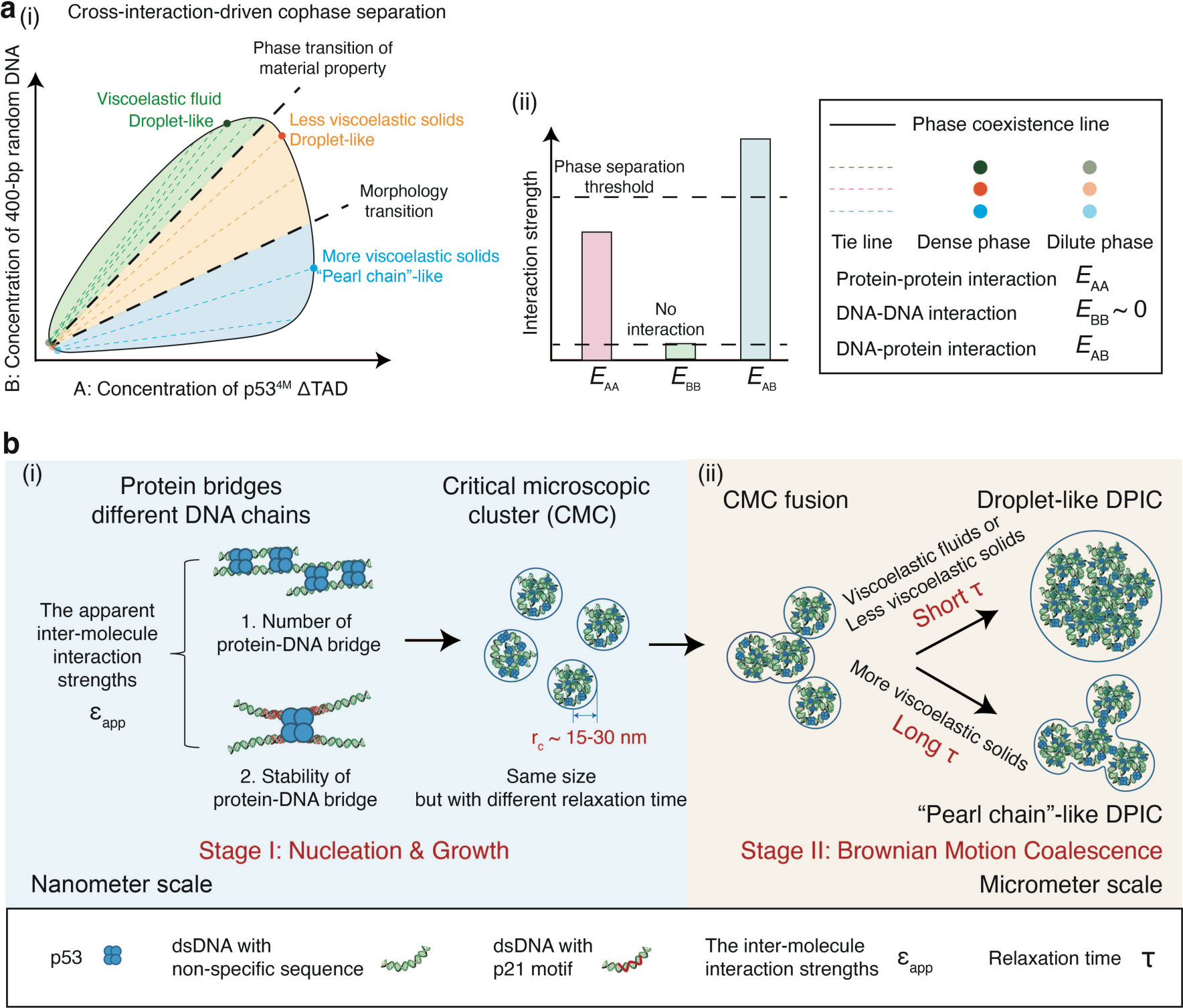
The molecular mechanism of DPICs. (**a**) (i) Schematic illustration of the two-component phase diagram of DPICs; (ii) Cross-interaction-driven phase separation: Component A (p53^4M^ ΔTAD) has some self-attraction (*E*_AA_), but cannot undergo liquid-liquid phase separation (LLPS), while component B (400-bp random DNA) has no attractive self-interaction (*E*_BB_ ∼ 0). (**b**) Schematic illustration of the growth dynamics of DPICs: (i) Stage I, nucleation and growth; (ii) Stage II, Brownian motion coalescence.

To validate this model, we drew upon established principles in soft matter physics concerning nucleation, growth, and coarsening (3, 57) for reference. Initially, we formulated a two-dimensional off-lattice model (Supplementary Fig. 9 & **Supplementary Methods**) to illustrate the regulatory role of ε_app_ in modulating relaxation time. Our analysis revealed that an increase in ε_app_ correlates with a decrease in the circularity of condensates (Supplementary Fig. 9b(iii)), accompanied by a prolonged relaxation time (Supplementary Fig. 9f(i)). Subsequently, we developed a mathematical model based on the viscoelastic phase separation (VPS) theory proposed by Hajime Tanaka (Supplementary Fig. 10 and **Supplementary Methods**) (58). Our inquiry aimed to ascertain whether manipulating the relaxation time parameter in this model could influence the morphology transition.

We optimized parameters such as the interaction parameter (χ), modulus of bulk stress (*G*_*B*_), modulus of shear stress (*G*_*S*_), and others, to fit the DPIC system (59). In the initial state, droplets of uniform size, which mimicked CMCs in the previous session, were randomly situated in a square box with periodic boundary conditions (Supplementary Fig. 10a(i), b(i), c(i), and d(i)). While maintaining all other parameters constant, we exclusively altered the maximum values of relaxation times (τ_max_) for bulk and shear stress, acting as regulators of the droplets’ mechanical properties. The systems evolved over 5,000 steps as per Eq. (1)-(3) in **Supplementary Methods**. The dynamics and final morphology of the condensates displayed substantial variability with τ_max_. A small τ_max_ (e.g., 0 or 1) led to rapid fusion, resulting in large, round droplets (Supplementary Fig. 10a(ii), b(ii), and Supplementary Movie 9-10). Conversely, a larger τ_max_ (e.g., 100) allowed for droplet fusion but maintained slower deformation, resulting in aspheric morphology (Supplementary Fig. 10c(ii) and Supplementary Movie 11). Extremely high τ_max_ (e.g., 10,000) led to the coalescence of initial droplets but hindered complete integration, preserving a “pearl chain”-like morphology (Supplementary Fig. 10d(ii) and Supplementary Movie 12). As τ_max_ was the only variable in this model, these outcomes underscored that altering relaxation time alone could be enough to induce morphological changes.

In summary, our investigation of the growth dynamics of DPICs bridges the microscopic parameter the ε_app_ with macroscopic morphology and material properties. We emphasize that interactions between DNA and protein initially induce CMCs at the nanometer scale. Variations in the ε_app_ result in distinct relaxation times, thereby dictating the diverse morphology and material properties of DPICs at the micrometer scale.

This enlightens us on whether it is possible to record the rounding of "pearl chain"-like DPIC by extending the observation time. The results showed that DPIC formed by 20 μM p53^4M^ ΔTAD and 0.6 μM 400-bp DNA containing 3× p21 binding motifs was “pearl chain”-like when the incubation time was only 30 minutes. Interestingly, when the incubation time was extended to 16 hours, the morphology of this sample changed greatly, with the circularity increased significantly (Supplementary Fig. 11). This indicates that “pearl chain”-like DPIC with higher ε_app_ does have a longer relaxation time, and it must be accompanied by a longer observation time to see the relaxation process.

## Discussion

NAPCs play a critical role in elucidating the complex physical chemistry underlying multi-component phase separations within living cells, consequently influencing various biological functions. Given that DPICs represent the simplest form of NAPCs, we employed p53 and dsDNA as a model system to investigate the biophysical mechanisms governing DPICs (Fig. 7). Our findings reveal that p53^4M^ ΔTAD can engage in DPIC formation with dsDNA, involving the bridging of distinct DNA duplexes by p53^4M^ ΔTAD (Fig. 7b(i)). The ε_app_ can be modulated by adjusting either the number of bridges on each DNA or the binding affinity between DNA and protein. To elucidate the molecular mechanisms underlying morphology transition, we initially delineated the phase diagram of DPICs (Fig. 7a(i)), which denotes a cross-interaction-driven cophase separation (Fig. 7a(ii)) (53). From this phase diagram, we identified two distinct transition behaviors: a phase transition between viscoelastic fluid and viscoelastic solid, and a morphology transition between droplet-like DPICs and those exhibiting a “pearl chain”-like morphology. Furthermore, we investigated the growth dynamics of DPICs. Interactions between DNA and protein led to the formation of CMCs with radii of approximately 15-30 nm. These CMCs, characterized by various viscoelastic properties, coalesce into larger DPICs with diverse morphologies.

Based on our data, several interesting topics warrant further discussion in future studies. First is about the two-component phase diagram of DPICs (Fig. 7a). Dignon and colleagues comprehensively reviewed four distinct types of phase diagrams for a two-component system (53): cooperative, scaffold-client, cross-interaction-driven, and exclusive cophase separation. Within the framework of cross-interaction-driven cophase separation, each species is incapable of undergoing phase separation independently, and only the mutual attraction between both species induces cophase separation. As discussed earlier, neither p53^4M^ ΔTAD nor dsDNA can undergo LLPS independently, suggesting that the protein-protein interactions (*E*_AA_) and the dsDNA-dsDNA interactions (*E*_BB_) are weak, with protein-dsDNA interactions (*E*_AB_) predominantly governing the overall DPIC system. Consequently, the phase paradigm characterizing our DPIC system, formed by the combination of p53^4M^ ΔTAD and 400-bp random DNA, represents a unique instance of the cross-interaction-driven cophase separation paradigm. Interestingly, in comparison to the cross-interaction-driven cophase separation model (*E*_AA_ = *E*_BB_ ∼ 0, and *E*_AB_ > 0) proposed by Dignon and colleagues (53), our phase diagram (Fig. 5c and Supplementary Fig. 8d) reveals dynamic changes in all positively sloped tie lines within the coexistence region (Fig. 7a(i)). We assumed that although p53^4M^ ΔTAD molecules are unable to undergo LLPS, there still exist inter-molecular interaction strengths between p53^4M^ ΔTAD molecules, indicating *E*_AA_ > 0 (Fig. 7a(ii)). This condition (*E*_AA_ > 0 but no LLPS, *E*_BB_ ∼ 0, and *E*_AB_ > 0) poses an interesting topic for future study.

We also notice that our cross-interaction-driven cophase separation paradigm delineates two distinct transition behaviors (Fig. 7a(i)). As the slope of tie lines progressively diminishes, the initial transition signifies a phase shift between viscoelastic fluid and viscoelastic solid. It is crucial to emphasize that the droplet-like morphology of DPICs persists during this phase transition. The second type of transition becomes evident as the slope of the tie lines continues to decrease, indicating a morphology transition from DPICs with droplet-like characteristics to those exhibiting a “pearl chain”-like morphology. These findings underscore that the phase transition from viscoelastic fluid to viscoelastic solid does not instantaneously alter the morphology of DPICs. Similar behavior was observed by Ephrussi and colleagues, who noted that oskar granules undergo a liquid-to-solid phase transition while maintaining a spherical shape (60); rather, the transition in morphology is distinctly driven by the presence of more viscoelastic solids. We speculate that altering the material properties of DPICs, transitioning between a viscoelastic fluid and a viscoelastic solid while maintaining the droplet-like morphology, could serve as a valuable strategy for living cells to finely regulate essential cellular processes. This includes processes like transcription and repair within various biomolecular condensates that involve nucleic acids. This is another interesting topic for the future study.

Second, drawing from established principles in soft matter physics concerning nucleation, growth, and coarsening (3, 57), in addition to BMC, the CMCs may undergo another process known as diffusion-limited coarsening (DLC) or Ostwald ripening (61, 62). This mechanism involves the dissolution of small condensates followed by the re-deposition of the dissolved molecules onto the surfaces of larger condensates, akin to the vapor pressure phenomena observed in soft matter physics (57). However, recent investigations led by Brangwynne and colleagues have highlighted a preference for biomolecular condensates to predominantly undergo coarsening via BMC (63). Our two-color experiments depicted in Fig. 1e provide additional support for BMC as the dominant mechanism during Stage II. Therefore, the exploration of whether biomolecular condensates can indeed undergo DLC remains an intriguing prospect for future inquiry.

Third, this conceptual framework of our model in Fig. 7 finds relevance in the context of a recent discourse on phase separation coupled to percolation (PSCP) (64). Through MD simulations (Fig. 2), we observed that DPICs exhibited an organized network structure attributed to the bridging effect of p53 between different DNA chains. Additionally, given the viscoelastic physical properties exhibited by DPICs (Fig. 4), we postulate that the formation of DPICs may entail a percolation process. This intriguing aspect warrants another further exploration in future studies.

Fourth is about reentrant phase behavior. Reentrant phase behavior characterizes the phenomenon wherein a change in a single thermodynamic parameter induces two or more phase transitions, resulting in the macroscopic state of the phase separation system becoming similar or identical to the original state (65). In the context of biomolecular condensates, two distinct reentrant phase behaviors have been observed. The first type involves a transition of “Two phases – One phase – Two phases”, which represents the phase region with an hourglass shape (66). For instance, at low or very high salt concentrations, proteins such as FUS, Sox2, and Brd4 can form condensates, but at a medium KCl concentration, these proteins can adopt a well-mixed state (67). The second type entails a transition of “One phase – Two phases – One phase”, as reported in some RNA-protein co-condensates involving arginine-rich peptides and homopolymeric single-stranded RNA, which represents the phase region with a closed loop (66). Condensates form only when the RNA concentration is at a medium level, with no condensates observed at low or high concentrations of RNA (13, 68). Our phase diagram in Fig. 1c indicates that DPICs also exhibit similar reentrant phase behavior. At 5 μM p53^4M^ ΔTAD, 0.15 μM 400-bp random DNA induces readily detectable droplet-like DPICs; however, at concentrations of 0.6 μM and above, droplet-like DPICs are no longer discernible. The detailed mechanism underlying reentrant phase behavior in DPICs warrants another further exploration in future studies.

In summary, our findings offer a quantitative characterization of the phase-separation mechanism of DPIC. Moreover, they present a novel avenue for comprehending the intricate world of multi-component condensates within living cells. Ultimately, these insights may pave the way for the manipulation of biomolecular condensation behaviors, offering innovative strategies for intervening in human diseases.

## Acknowledgments

We thank the Peking Nanofab for process support. We thank the contributions of the Engineering Research Center for Semiconductor Integrated Technology, Institute of Semiconductors, Chinese Academy of Sciences. We thank the National Center for Protein Sciences at Peking University in Beijing, China, particularly Dr. Siying Qin, for technical help with AFM. We thank the computational support from the High-Performance Computing Center of Nanjing University. We thank Dr. Luhua Lai (Peking University), Dr. Sheng Mao (Peking University), Dr. Ming Han (Peking University), Dr. Chun Tang (Peking University), and members of the Qi laboratories for comments on the manuscript.

## Author Contributions

C.L. prepared biological samples, conducted all experiments, performed data analysis, and wrote the manuscript. Y.B. and W.L. conducted the coarse-grained molecular dynamics and two-dimensional off-lattice model. Y.T. and C.C. conducted the dcFCCS experiments and performed data analysis. L.M. and J.L. conducted numerical simulations of viscoelastic phase separation. P.Y. developed *in vitro* p53 purification protocol and *in vitro* droplet assay for p53 condensate study. Y.H. and J.C. assisted C.L. Y.L. helped to purify p53^4M^ ΔTAD. Z.Q., W.L., and C.T. supervised the project, experimental designs, and data analysis. Z.Q. and W.L. wrote the manuscript with input from all authors.

## Methods

### Construction of bacterial expression plasmids

It has been previously proved (36) that four mutations (M133L/V203A/N239Y/N268D) on wild type (WT) human p53 could make the protein stable *in vitro*. We cloned full-length p53 on pRSFDuet vector, mutated those bases and introduced His-tag to construct the plasmid (No. 1) – 6×His-p53^4M^. From N-terminal to C-terminal, p53^4M^ can be divided into following domains: NTD (1-94), DBD (95-292), TET (293-356) and CTD (357-393). NTD and CTD are N-terminal domain and C-terminal domain, DBD is the middle core domain that is also DNA binding domain, and TET is the tetramerization domain. NTD includes two TADs (tandem transcription activation domain) and one PRD (Proline-rich domain). We deleted the TAD1 and TAD2 to get a new plasmid (No. 2) – 6×His-PRD-DBD-TET-CTD (p53^4M^ ΔTAD). Protein sequences were shown in the excel below.

### Protein purification and labeling

The expression plasmids for all proteins above were transformed into *E. coli* strain BL21(DE3), and then cultured overnight on LB agar plates. A single clone obtained from the plate was cultured in 10 mL LB medium overnight at 37 °C and 220 rpm with 50 μg/mL Kanamycin. The culture was diluted by 1:250 into 2 L LB medium, and then grew to a density (OD600 nm) of 0.6. Bacteria were induced with 0.3 mM IPTG and 0.1 mM ZnCl_2_ at 16 °C and 180 rpm overnight. Cultures were centrifuged with 4,000 g. Cells were resuspended in a lysis buffer (25 mM Tris-HCl (pH 7.5), 500 mM NaCl, 5 mM imidazole, 0.25‰ β-Mercaptoethanol (β-Me), and 5% Glycerol), and then sonicated with 1 mM PMSF on ice. Supernatant was collected and filtered after centrifugation at 18,000 rpm for 40 minutes. Ni-NTA was used to bind p53, and the beads were washed by a washing buffer (25 mM Tris-HCl (pH 7.5), 500 mM NaCl, 20 mM imidazole, 0.25‰ β-Me, and 5% Glycerol). Finally, the protein was eluted by an elution buffer (25 mM Tris-HCl (pH 7.5), 500 mM NaCl, 300 mM imidazole, 0.25‰ β-Me, and 5% Glycerol), and stored at -80 °C in a storage buffer (20 mM Tris-HCl (pH 7.5), 300 mM NaCl, 10% Glycerol, and 40 mM DTT) after further purification by gel filtration with a Superdex 200 increase GL 10/300 (GE Healthcare, USA). Purified proteins were checked by SDS-PAGE.

In the FRAP assay, p53^4M^ ΔTAD was labeled by ATTO565 NHS ester (Sigma Cat: 72464) at 1:1.5 molar ratios in 0.1 M NaHCO_3_ for 1 hour at 4 °C. In the dcFCCS assay, p53^4M^ ΔTAD was labeled by Sulfo Cyanine5 (Cy5) NHS ester (Lumiprobe) at 1:4 molar ratios in reaction buffer (20 mM HEPES pH 7.5, 150 mM KCl, 10% glycerol, 1 mM TCEP) and incubated for 20 min at RT. After labeling, the labeled protein molecules were separated from the free dye and changed into storage buffer (20 mM Tris-HCl (pH 7.5), 300 mM NaCl, 10% Glycerol, and 40 mM DTT) on the Superdex 200 increase GL 10/300. Then labeled protein samples were stored at -80 °C.

### Electrophoretic Mobility Shift Assay (EMSA)

#### EMSA

EMSA has been performed to test the binding affinity of proteins used in this paper on DNA. 199-bp DNA probe with or without p53 specific binding site was amplified from plasmid by PCR with one primer labeled by 5’-quasar 670 and prepared at a final concentration of 500nM in ddH_2_O. The DNA sequences are shown in the excel below. The DNA binding reaction was performed with a DNA probe of 5 nM final concentration in the p53 working buffer (20 mM HEPES (pH 7.9), 150 mM NaCl, 2 mM MgCl_2_, 1 mM DTT, and 0.5 mg/mL BSA). p53^4M^ or truncated version protein was serial diluted and mixed with DNA probe at different working concentration. DNA-protein mixture was incubated for 30 minutes at room temperature (RT) and resolved by 1% agarose gel in TBE buffer under 80 V for 80 minutes in cold room (4 °C). The results of EMSA are identified from scanning through the Cy5 channel by Amersham Typhoon.

### *In vitro* droplet assay

#### *In vitro* droplet assays

All *in vitro* samples were dropped into 384-well plates (Cellvis). If not mentioned, the p53 droplet buffer was 8 mM Tris-HCl (pH 7.5), 120 mM NaCl, 4% Glycerol, and 16 mM DTT. The salt concentration was changed in different experiments. The p53^4M^ or truncated version protein and DNA with different length or sequence are centrifuged at 12,000× g for 10 min at 4 °C before usage, and then diluted to different working concentration by storage buffer or ddH_2_O. The DNA-protein mixture is incubated at room temperature (RT) for 0.5 hour. Confocal (Leica TCS SP8) was used to observe DPIC.

#### Fluorescence recovery after photobleaching (FRAP)

For the FRAP experiment, the DNA-protein mixture was incubated at RT for 2 hours to obtain sufficiently large droplet-like condensates. We used a spinning-disk confocal microscopy (UltraView VoX) to bleach an area of 1.5 μm × 1.5 μm inside each droplet by 488 nm and 561 nm laser with strength of 100% for 30 cycles, and recorded the recovery process with a speed of 2 minutes per frame for 120 minutes. With the program described above, we obtained a chronological series of photos, and analyzed fluorescence intensity of the bleached regions together with controlled ones to normalize and plot a curve (Supplementary Fig 3).

#### Fluorescence intensity measurement of protein and DNA inside DPIC

The DPICs formed by ATTO565 labeled p53^4M^ ΔTAD and FAM labeled 400-bp random DNA were incubated at RT for 30 min, and then scanned by 488- and 552-nm lasers on Leica TCS SP8 microscope. The laser strength and the gain value of detectors were fixed during observation. Images were analyzed with Fiji (ImageJ Version: 2.0.0-rc-61/1.52n). The image in fluorescent channel was first adjusted to 8-bit. For each field of observation, 10 DPICs were selected, and the average fluorescence intensity per pixel inside them was measured in ATTO565 channel and FAM channel, respectively. Three independent fields of view were measured for each experimental condition.

#### Droplet circularity analysis

To obtain the circularity of condensates, the image in fluorescent channel was first adjusted to 8-bit. Threshold of the image was automatically set to identify particles with the subroutine of “threshold” in Fiji (Image/Adjust/Threshold). Particles larger than 1 μm^2^ were selected for shape descriptors analysis with the subroutine of “analyze particles” in Fiji (Analyze/Analyze particles). The average circularity can be calculated from all measured values.

To measure the circularity of DPICs formed by the two-dimensional off-lattice model, the snapshots of all condensates at 5× 10^8^ MD steps were first binarized with the subroutine of “Make binary” in Fiji (Process/Binary/Make binary). To reduce the effect of rough boundaries caused by simplified structure of p53 tetramer and dsDNA, the images were undergo smoothed with the subroutine of “Smooth” in Fiji (Process/Smooth). Threshold of the images was set automatically, and the condensates larger than 100 pixels were measured.

### Dual-color Fluorescence Cross-correlation Spectroscopy (dcFCCS)

The dcFCCS measurements were performed by a home-made confocal microscope with 488- and 640-nm lasers. More details about the setup of microscope can be referred to the previous procedure (55). Powers of lasers were set about 5 μW after the objective. The laser focus was 10 μm above the coverslip-water interface. The temperature was 25 °C. In all dcFCCS measurements, the working buffer was 8 mM Tris-HCl (pH 7.5), 120 mM NaCl, and 16 mM DTT. It took 5 min to mix protein and DNA well, to load them into the reaction chamber, to turn on laser and detectors, and to start collecting data. Raw data of photon arriving time of FAM detection channel (ET525/50m, Chroma) and Cy5 detection channel (ET700/75m, Chroma) was recorded for 5 min. For each experimental condition, at least 5 repeats were performed. Auto-correlation curves of FAM detection channel and Cy5 detection channel and cross-correlation curve between them were calculated from the raw photon arriving time using a home-made MATLAB script.

### Atomic Force Microscopy-based Force Spectroscopy (AFM-FS)

For AFM-FS experiments, all DPIC samples were incubated at RT for 1 hour and washed by the droplet buffer, removing all DPICs which cannot adhere tightly to the coverslip surface. Measurements were performed with the operation mode of force volume mode in fluid on the commercial AFM BioScope Resolve (Bruker, Billerica, MA, USA). A silicon nitride probe (PFQNM-LC-A-CAL, Bruker) with a tip height of 17 μm, a tip radius of 70 nm, and pre-calibrated spring constant of 0.076 N/m was used for the force measurements. All the force curves were recorded with the loading and unloading speed of 2 μm/s, scan size of 50 nm, ramps/line of 4, ramp size of 2 μm, ramp rate of 0.5 Hz, and the deflection error triggers a threshold of 13 nm. For the stress relaxation test, the tip was hold at a fixed height above the DPIC for 1 second by using the hold type of Z-drive after the deflection error trigger threshold was reached. The sampling rate is 1,024 Hz. All experiments were done at RT within 1 hour. Each DPIC was indented for 16 times and about 20 different DPICs were tested for each condition.

All AFM results were imported into the Nanoscope Analysis software (Bruker). For each DPIC, according to whether the force baselines of extend or retract overlap, three sets of force curves (n = 3) with good overlap were picked from 16 sets of stress relaxation test manually. The values of force baseline of picked curves were then corrected to 0 through the Baseline correction of the software with 0^th^ correction order. The force-time, height-time and separation-time curves of each DPIC were derived. The force-time curves were then imported into the Excel software. The time range of the hold process was picked out. To calculate the normalized force, the force value of each time point was divided by the maximum value during the hold process.

### Unpaired t test

Statistical significance was evaluated based on Student’s t-tests (Prism 9 for macOS, Version 9.1.0 (216), March 15, 2021, GraphPad Software, Inc.). Test was chosen as unpaired t test. P value style: GP: 0.1234 (ns), 0.0332 (*), 0.0021 (**), 0.0002 (***), < 0.0001 (****).

### Box-plot

The function of “boxplot” in MATLAB software (R2015a, 64-bit, February 12, 2015) was used to plot the boxplots in Fig. 3-4 and Supplementary Fig. 7 and 11. For each boxplot, the red bar represents median. The bottom edge of the box represents 25^th^ percentiles, and the top is 75^th^ percentiles. Most extreme data points are covered by the whiskers except outliers. The ‘+’ symbol is used to represent the outliers.

