## Supplemental files for "Deciphering the molecular mechanism underlying morphology transition in two-component DNA-protein cophase separation"

**Table of contents**

1. **Supplementary Methods**
   - **1.1 Coarse-grained MD simulation**
   - **1.2 A two-dimensional off-lattice model**
   - **1.3 Numerical simulations of viscoelastic phase separation**
   - **1.4 Generated Maxwell model**
2. **Oligonucleotide preparation**
3. **Protein sequences**
4. **Supplementary Figures**
5. **Supplementary Movie Legends**

### 1. Supplementary Methods

#### 1.1 Coarse-grained MD simulation

The molecular simulations were performed using a coarse-gained model. For protein, each residue was represented by a single particle centered on its C$\alpha$ position (Supplementary Fig. 4a). The AICG2+ energy function, which consists of a flexible local potential term, a structure-based potential term, and an excluded volume term, was used to describe the interactions and motions of the folded domains of p53$^{4M}$ (41, 42). Meanwhile, the HPS energy function, which describes the generic hydrophobic interactions for polypeptide chains, was employed to characterize the interactions and motions of intrinsic disordered regions (IDRs) in p53$^{4M}$ (69, 70). For DNA, each nucleotide is represented by three beads corresponding to phosphate, sugar, and base, respectively. The 3SPN.2C energy function developed by de Pablo's group was used (39). In addition, the Debye-Hückel type electrostatic potential was used to characterize the salt-concentration-dependent interactions of charged particles in protein and DNA. In describing the interactions between specific DNA sequences (1$\times$ p21 binding motif) and protein, the PWMcos model developed by Tan and Takada, which integrates high-throughput protein binding assays of transcription factors and structural biology experiments (71), was used. Details of the coarse-grained model can be found elsewhere (39, 42, 69). The combination of AICG2+ and 3SPN.2C models has shown great success in the simulations of various molecular processes, such as the p53 sliding on DNA chain, the nucleosome remodeling and allostery, and the dynamic coupling of protein binding and DNA bending (40, 72, 73).

All simulations were conducted with the GENESIS 1.7.1 package (44-46). The reference structures for constructing the intra-domain interactions of the folded domains and the inter-domain interactions between the OD domains in $p53^{4M}$ were taken from the Protein Data Bank with the entries of 2XWR and 1AIE. For the $p53^{4M}$ ΔTAD, the N-terminal 61 residues corresponding to the negatively charged TAD domain were truncated. The initial structures of all the DNA were built by the CG-tool packaged in GENESIS. Each system was placed in a cubic box with periodic boundary condition. The simulations were performed in NVT ensemble at a temperature of 300 K controlled by the Langevin dynamics with an inverse friction constant of 0.1 ps and the timestep of 0.01 ps. The salt concentration was set to 150 mM except otherwise stated. We conducted the simulations for the system with two different sizes. For the larger system, it includes seven dsDNA chains with a length of 199-bp and 27 $p53^{4M}$ ΔTAD tetramers, which were contained in a cubic box with the size of 750Å × 750Å × 750Å. The corresponding concentration is higher than the average concentration used in experiment. Two independent simulations lasting for $1 \times 10^8$ MD steps were conducted. To achieve improved sampling and more quantitatively characterize the effects of different factors on the capability of each $p53^{4M}$ tetramer to bridge multiple dsDNA chains, we also conducted the simulations for a much smaller system with the cubic box size of 550Å × 550Å × 550Å. It contains two $p53^{4M}$ ΔTAD (or full-length $p53^{4M}$) tetramers and 10 dsDNA with the length of 199 bp (or 30 bp). The total simulation time of each trajectory was $1 \times 10^8$ MD steps. Three independent simulations were conducted for each system.

In characterizing the capability for each $p53^{4M}$ tetramer to bridge multiple dsDNA chains, we used the collective variable $N_{DNA}$, i.e., the number of DNA chains bound by each p53$^{4M}$ tetramer. In order to calculate $N_{DNA}$, we first defined a variable $N_{ij}$, which describes the number of formed contacts between a DNA chain i and a p53$^{4M}$ monomer j. In this work, the residue bead and nucleotide bead form a contact when their distance becomes less than 6.5 Å (Supplementary Fig. 4a). A DNA chain was thought to bind with a p53$^{4M}$ monomer when the contact number between them is larger than a threshold of 5, by which we can calculate the collective variable $N_{DNA}$ and its distribution. We also tested the threshold values of 10 and 15, and similar results were obtained (Supplementary Fig. 5c). In calculating the Lifetime of the p53 core DBD binding with one DNA duplex, the p53 core DBD was considered as fully detached from the dsDNA when the contact number becomes zero. The Lifetime was estimated based on all the unbinding events collected from the three independent simulations at each case.

The above smaller system represents a minimal system for a quantitative discussion. As a sensitivity test of the results on the included number of p53$^{4M}$ and dsDNA chains, we also examined other small systems: (i) 2 p53$^{4M}$ ΔTAD tetramers and 20 199 bp dsDNA chains (Supplementary Fig. 5b(i)), $N_{DNA}$ = 3.4 ± 0.7 (n = 3); (ii) 5 p53$^{4M}$ ΔTAD tetramers and 10 199 bp dsDNA chains (Supplementary Fig. 5b(ii)), $N_{DNA}$ = 3.8 ± 0.6 (n = 3); (iii) 10 p53$^{4M}$ ΔTAD tetramers and 10 199 bp dsDNA chains (Supplementary Fig. 5b(iii)), $N_{DNA}$ = 3.5 ± 0.9 (n = 3). The results showed that the calculated $N_{DNA}$ values are comparable to that of the above minimal system containing 2 p53$^{4M}$ ΔTAD tetramers and 10 random dsDNA with the length of 199 bp (i.e., $N_{DNA}$ = 3.5 ± 0.7 in Supplementary Fig. 5a(i)). For the sake of computational feasibility, we used the above minimal system in the quantitative discussion of the effects of different factors on the capability of each p53$^{4M}$ ΔTAD to bridge multiple dsDNA chains.

### 1.2 A two-dimensional off-lattice model

To investigate the general physical characteristics of phase separation, a two-dimensional simplified model was constructed. In this model, p53 tetramer was modelled by a star-like structure with five beads. The four p53 monomers were represented by four arm beads, which were connected by a center bead, forming a tetramer structure (Supplementary Fig. 9a(i)). Similarly, the dsDNA was modelled as a linear chain consisting of seven beads. Each bead roughly represents 20 base pairs. The bonds between the successive beads were described by a spring potential $V_b(\theta) = k_b(r - r_b)^2$, with $r_b$ and $k_b$ being set as 1.2σ and 13500 kJ/mol/$\sigma^2$, respectively. Here σ represents the reduced unit of length, and *r* is the distance between two beads. Similarly, the bond angle formed by the neighboring three beads were applied a spring potential $V_\theta(\theta) = k_\theta(\theta - \theta_0)^2$, with $\theta$ and $\theta_0$ being the angle and its equilibrium value. In this work, $\theta_0$ was set to $\pi/4$ and $\pi$, respectively, for the p53 tetramer and dsDNA. $k_\theta$ was set to 10000 kJ/mol/$rad^2$. The interactions between the dsDNA beads and the p53-tetramer beads were modeled using

$$U_{LJ} = 4\lambda\varepsilon_0\left[\left(\frac{r_0}{r}\right)^{12} - \left(\frac{r_0}{r}\right)^{6}\right]$$

In the above formula, $r_0$ represents the bead size and was set as 1.0σ. $\varepsilon_0$ represents the energy scale and was set as 1.0kJ/mol. The coefficient $\lambda$ controls the interaction strength between p53 and dsDNA, which determines the structural and dynamic behavior of the system. By changing the coefficient $\lambda$, we can investigate the dependence of the phase separation behavior on the inter-molecule interaction strength. The coarse-grained beads within the same molecules were further applied an excluded volume interaction term. In this work, we performed the simulations using Langevin dynamics for a system containing 2,400 p53 tetramers and 1,400 dsDNA chains. These molecules were randomly placed into a square box with the size of ~$333\sigma \times 333\sigma$. Periodic boundary condition was applied in the simulations. The temperature was set to 298K. Each simulation lasted for a total of $5 \times 10^8$ MD steps.

#### 1.3 Increasing the $\varepsilon_{app}$ leads to a prolonged relaxation time

We developed a two-dimensional off-lattice model (**Supplementary Methods 1.2**), wherein $p53^{4M}$ ΔTAD was represented as a star-shaped structure comprising four branching beads linked to a central bead, while DNA was depicted as a straight but flexible chain composed of seven beads (Supplementary Fig. 9a(i)). The branching beads of $p53^{4M}$ ΔTAD tetramer interact with the DNA beads through the Lennard-Jones potential with the strength controlled by a parameter $\lambda$ (Supplementary Fig. 9a(ii)). The simulation system contains 2,400 $p53^{4M}$ ΔTAD tetramers and 1,400 DNA chains, which were randomly placed in a squared box of ~333 $\sigma \times$ 333 $\sigma$, with $\sigma$ being the reduced length unit. We performed Langevin dynamics simulations at the temperature of 298 K. When the $\lambda$ values are larger than 10, the molecules in the system formed large condensates, which allows us to investigate the relation between the $\varepsilon_{app}$ and the physical properties of the formed condensates.

As shown by the final structures of the simulations, the condensates were nearly round-shaped when $\lambda$ was low, suggesting liquid-droplet like property (Supplementary Fig. 9b(i) and Supplementary Movie 7). In comparison, the condensates became "pearl chain"-like when $\lambda$ was higher (Supplementary Fig. 9b(ii) and Supplementary Movie 8).

More quantitative characterization showed that increasing interaction strength $\lambda$ tends to decrease the circularity of condensates (Supplementary Fig. 9b(iii)). Such results were highly consistent with the above experimental observation that increased protein-DNA interaction strengths tend to drive the formation of condensates with "pearl chain"-like morphology. We also characterized the dynamic property of the condensate by calculating the diffusion coefficient (D) of the protein molecules within the condensates (Supplementary Fig. 9c-e). As expected, the diffusion coefficient decreases monotonically with the increasing of protein-DNA interaction strengths, demonstrating arrested molecular mobility with stronger protein-DNA interactions. We further calculated the diffusion time ($\tau_D = L^2/4D$, with L being the radius of the observed droplet) to roughly characterize the relaxation time ($\tau$) of protein-DNA scaffold inside the DPICs, which showed an anticorrelation with the $\varepsilon_{app}$ (Supplementary Fig. 9f(i)) and an anticorrelation with circularity (Supplementary Fig. 9f(ii)).

#### 1.4 Numerical simulations of viscoelastic phase separation

Numerical simulations of viscoelastic phase separation are based on the two-fluid model (74, 75), which considers the dynamics of biomolecule velocity $v_p$, solvent velocity $v_s$ and the average velocity $v = \phi v_p + (1-\phi)v_s$. In the two-fluid model, the biomolecule density field $\phi$ is spatially dependent with its value between 0 and 1. The dynamics of the biomolecule density and velocity field follows

$$\frac{\partial \phi}{\partial t} = -\nabla \cdot (\phi v_p), \tag{1}$$

$$v_p - v = -\frac{(1-\phi)^2}{\zeta}(\nabla \cdot \Pi - \nabla \cdot \sigma), \tag{2}$$

$$-\nabla \cdot \Pi + \nabla \cdot \sigma - \nabla p + \eta \nabla^2 v = 0. \tag{3}$$

Here, $\zeta$ is the friction constant between biomolecules and solvent, and $\eta$ is the viscosity. The pressure $p$ is determined by the incompressible condition: $\nabla \cdot v = 0$. Combined with Eq. (3), the average velocity $v$ can be solved as

$$\boldsymbol{v}(\boldsymbol{r}) = \int d\boldsymbol{r}' \, \boldsymbol{T}(\boldsymbol{r} - \boldsymbol{r}') \cdot \left(-\nabla \cdot \boldsymbol{\Pi}(\boldsymbol{r}') + \nabla \cdot \sigma(\boldsymbol{r}')\right), \tag{4}$$

while $T(k) = \frac{1}{\eta |k|^2}\left(I - \frac{kk}{|k|^2}\right)$ is the Oseen tensor in the Fourier space. We can then obtain the biomolecule velocity $v_p$ with Eq. (2). Therefore, the density $\phi$ in simulation can be updated with Eq. (1) by calculating $v_p$ when the osmotic pressure $\Pi$ and the stress $\sigma$ are known. The osmotic stress tensor is determined by the biomolecule free energy $f(\phi)$, $\nabla \cdot \Pi = \phi \nabla f'(\phi)$, where we use the Flory-Huggins free energy density $f(\phi) = \epsilon_0 \left[\phi \ln(\phi) + (1-\phi)\ln(1-\phi) + \chi\phi(1-\phi) + \frac{C}{2}(\nabla\phi)^2\right]$ and $C$ is a constant. Here $\epsilon_0 = k_B T / V_0$ where $k_B$ is the Boltzmann constant, $T$ is the temperature, and $V_0$ is the biomolecular monomer volume. The stress tensor$\sigma = \sigma_S + \sigma_B I$ where $\sigma_B$ is the bulk stress and $\sigma_S$ is the shear stress tensor. Based on the experimental results (59), they follow the Maxwellian dynamics such that

$$\frac{\partial \sigma_B}{\partial t} = -\left(v_p \cdot \nabla\right)\sigma_B - \frac{1}{\tau_B(\phi)}\sigma_B + G_B(\phi)\nabla \cdot v_p, \tag{5}$$

$$\begin{aligned} \frac{\partial \sigma_S}{\partial t} &= -\left(v_p \cdot \nabla\right)\sigma_S + \sigma_S \cdot \nabla v_p + \left(\nabla v_p\right)^T \cdot \sigma_S \\ &\quad - \frac{1}{\tau_S(\phi)}\sigma_S + G_S(\phi)\left(\nabla v_p + \left(\nabla v_p\right)^T\right). \end{aligned} \tag{6}$$

Here $G_B(\phi)$ is the bulk modulus and $G_S(\phi)$ is the shear modulus. In the numerical simulations, we take

$$G_B(\phi) = G_B \Theta(\phi - \phi_c), \qquad G_S(\phi) = G_S \phi^2, \tag{7}$$

where $G_B$ and $G_S$ are constants and $\Theta(x)$ is the Heaviside function. We assume a critical density $\phi_c$ above which the biomolecule network is percolated and becomes elastic with a finite bulk modulus and relatively large relaxation time as

$$\tau_B(\phi) = \tau_S(\phi) = 0.01 + \frac{\tau_{max}}{2}\left[1 + \tanh\left(100(\phi - \phi_c)\right)\right]. \quad (8)$$

By non-dimensionalizing Eqs. (1-3), we choose the unit of elastic modulus as $\epsilon_0$, the time unit $t_0 = \eta/\epsilon_0$, and the length unit $l_0 = \sqrt{\eta/\zeta}$. We perform numerical simulations in a 2D 511*511 grid by solving the two-fluid model using the explicit Euler method with the periodic boundary condition on MATLAB. Eq. (4) is solved with fast Fourier transformation, and other equations are calculated in real space. For the Flory-Huggins free energy, we choose $\chi = 3$. For the Maxwellian dynamics, we choose $G_B = G_S = 10$ and $\phi_c = 0.7$. In our simulations, we take $C = 1$ and the grid size is $\Delta l = 0.25$. The total time for the simulation is $t_{\mathrm{tot}} = 5000$ and the time interval for the simulation is $\Delta t = 0.005$. For the initial condition, we generate 265 condensates with initial radius $R_{\mathrm{ini}} = 4$, where the inside and outside volume fractions of the condensates are the equilibrium volume fractions at $\chi = 3$ respectively.

#### 1.5 Generated Maxwell model

This non-covalent crosslinked structure is generally considered to exhibit physical characteristics of viscoelasticity (76). Unlike the ideal elastomers (all the deformation energy done by the external force is stored without any loss, that is, the elastic response) or the ideal viscous fluid (the deformation energy done by the external force is completely lost when it flows, which cannot store any energy, that is, the viscous response), viscoelastic materials can exhibit both elastic response and viscous response, and can behave hysteresis, stress relaxation, creep and other properties in response to input force (50, 77).

To quantitatively measure the viscoelasticity, we conducted the AFM experiment: stress relaxation test (Fig. 4a-b) (50). A generated Maxwell model consisting of multiple parallelly arranged Maxwell elements was used to fit the stress relaxation curve (51, 52, 78, 79). The formula for the model is $\mathrm{F(t)} = A_0 + \sum_{i=1}^{N} A_i \exp\left(-\frac{t}{\tau_i}\right)$. In this formula, $A_0$ is an instantaneous response and can be considered as the purely elastic effect. $A_i$ is the force decay and $\tau_i$ is the relaxation time of each Maxwell element. A cartoon figure in Fig. 4b(iii) was used to explain the generated Maxwell model used in this work, which consists of a spring and two Maxwell elements in parallel. After the stress relaxation curves were normalized (Fig. 4b), the original formula can be rewritten as $\mathrm{F(t)} = (1 - A_1 - A_2) + A_1 \exp\left(-\frac{t}{\tau_1}\right) + A_2 \exp\left(-\frac{t}{\tau_2}\right)$. $\tau_1$ and $\tau_2$ correspond to the relaxation time and $A_1$ and $A_2$ correspond to the force decay of the two Maxwell element. So, the force remain can be calculated as $A_0$ (= 1 – $A_1$ – $A_2$) to represent the purely elasticity of DPIC. When $A_0 = 0$, the DPIC is viscoelastic fluid. When $A_0 > 0$, the DPIC is viscoelastic solid (50).

### 2. Oligonucleotide preparation

We ordered 5× p21 sequence from Ruibiotech. Between each two p21 target sequences was a 17-bp linker. The sequence was cloned into pUC57 vector to get (No. 3) – 5× p21 plasmid. Non-specific DNA (199-bp) was a 199 bp sequence from the vector with no p21 target in it. All DNA samples used in our *in vitro* droplet experiments and EMSAs were prepared through PCR from lambda DNA and plasmid above, then purified by the spin column (FastPure Gel DNA Extraction Mini Kit, DC301-01, Vazyme). FAM labeled dsDNA was prepared by PCR with one primer labeled by 5'-FAM. DAPI labeled dsDNA was prepared by addition of 1 mg/mL DAPI (DAPI, D9542, Sigma-Aldrich) to DNA stock at a volume ratio of 1:1,000.

30-bp and 60-bp DNA substrates used for *in vitro* droplet experiments and EMSAs were generated by annealing. In the annealing system, top strands and bottom strands were added followed the molar ratio of 1:1.2 in annealing buffer containing 40 mM Tris-HCl (pH 8.0), 50 mM NaCl, and 10 mM $MgCl_2$. Then the tube was heated with a heater at 95 $^{o}$C for 5 minutes, and slowly cooled down to room temperature over 40 minutes together with the heater.

Oligonucleotide sequences we used are shown below. Relevant target sequences are emphasized with underlines:

| Oligo application | Sequence (5' to 3'), p21 sequence shown in red |
|---|---|
| 30-bp random DNA for EMSA | AAC CTG TCG TGC CAG CTG CAT TAA TGA ATC |

| | |
|---|---|
| 30-bp DNA with 1× p21 binding motif for EMSA | TCT AT GAA CAT GTC CCA ACA TGT TG TTC CC |
| 199-bp random DNA for EMSA and *in vitro* droplet assay | AGT GAG CTA ACT CAC ATT AAT TGC GTT GCG CTC ACT GCC CGC TTT CCA GTC GGG AAA CCT GTC GTG CCA GCT GCA TTA ATG AAT CGG CCA ACG CGC GGG GAG AGG CGG TTT GCG TAT TGG GCG CAC TAC CGC TTC CTC GCT CAC TGA CTC GCT GCG CTC GGT CGT TCG GCT GCG GCG AGC GGT ATC AGC TCA CTC AAA G |
| 199-bp DNA with 5× p21 binding motifs for EMSA | TGA ATT CCT CGA GGA ACA TGT CCC AAC ATG TTG AGC TCT GGC ATA GAA GAG AAC ATG TCC CAA CAT GTT GAG CTC TGG CAT AGA AGA GAA CAT GTC CCA ACA TGT TGA GCT CTG GCA TAG AAG AGA ACA TGT CCC AAC ATG TTG AGC TCT GGC ATA GAA GAG AAC ATG TCC CAA CAT GTT GCT AGC AAG CTT GGC GTA A |
| 30-bp random DNA for *in vitro* droplet assay | AAC CTG TCG TGC CAG CTG CAT TAA TGA ATC |
| 60-bp random DNA for *in vitro* droplet assay | TTG CGC TCA CTG CCC GCT TTC CAG TCG GGA AAC CTG TCG TGC CAG CTG CAT TAA TGA ATC |
| 120-bp random DNA for *in vitro* droplet assay | CTT TTG CTT GAT CTC AGT TTC AGT ATT AAT ATC CAT TTT TTA TAA GCG TCG ACG GCT TCA CGA AAC ATC TTT TCA TCG CCA ATA AAA GTG GCG ATA GTG AAT TTA GTC TGG ATA GCC ATA |

| | |
|---|---|
| 400-bp random DNA for *in vitro* droplet assay | CTT TTG CTT GAT CTC AGT TTC AGT ATT AAT ATC CAT TTT TTA TAA GCG TCG ACG GCT TCA CGA AAC ATC TTT TCA TCG CCA ATA AAA GTG GCG ATA GTG AAT TTA GTC TGG ATA GCC ATA AGT GTT TGA TCC ATT CTT TGG GAC TCC TGG CTG ATT AAG TAT GTC GAT AAG GCG TTT CCA TCC GTC ACG TAA TTT ACG GGT GAT TCG TTC AAG TAA AGA TTC GGA AGG GCA GCC AGC AAC AGG CCA CCC TGC AAT GGC ATA TTG CAT GGT GTG CTC CTT ATT TAT ACA TAA CGA AAA ACG CCT CGA GTG AAG CGT TAT TGG TAT GCG GTA AAA CCG CAC TCA GGC GGC CTT GAT AGT CAT ATC ATC TGA ATC AAA TAT TCC TGA TGT ATC GAT ATC GGT A |
| 600-bp random DNA for *in vitro* droplet assay | CTT TTG CTT GAT CTC AGT TTC AGT ATT AAT ATC CAT TTT TTA TAA GCG TCG ACG GCT TCA CGA AAC ATC TTT TCA TCG CCA ATA AAA GTG GCG ATA GTG AAT TTA GTC TGG ATA GCC ATA AGT GTT TGA TCC ATT CTT TGG GAC TCC TGG CTG ATT AAG TAT GTC GAT AAG GCG TTT CCA TCC GTC ACG TAA TTT ACG GGT GAT TCG TTC AAG TAA AGA TTC GGA AGG GCA GCC AGC AAC AGG CCA CCC TGC AAT GGC ATA TTG CAT GGT GTG CTC CTT ATT TAT ACA TAA CGA AAA ACG CCT CGA GTG AAG CGT TAT TGG TAT GCG GTA AAA CCG CAC TCA GGC GGC CTT GAT AGT CAT ATC ATC TGA ATC AAA TAT TCC TGA TGT ATC GAT ATC GGT AAT TCT TAT TCC TTC GCT ACC ATC CAT TGG AGG CCA TCC TTC CTG ACC ATT TCC ATC ATT CCA GTC GAA CTC ACA CAC AAC ACC ATA TGC ATT TAA GTC GCT TGA AAT TGC TAT AAG CAG AGC ATG TTG CGC CAG CAT |

| | |
|---|---|
| | GAT TAA TAC AGC ATT TAA TAC AGA GCC GTG TTT ATT GAG TCG GTA TTC AGA GTC TGA CCA GAA |
| 774-bp random DNA for *in vitro* droplet assay | CTT TTG CTT GAT CTC AGT TTC AGT ATT AAT ATC CAT TTT TTA TAA GCG TCG ACG GCT TCA CGA AAC ATC TTT TCA TCG CCA ATA AAA GTG GCG ATA GTG AAT TTA GTC TGG ATA GCC ATA AGT GTT TGA TCC ATT CTT TGG GAC TCC TGG CTG ATT AAG TAT GTC GAT AAG GCG TTT CCA TCC GTC ACG TAA TTT ACG GGT GAT TCG TTC AAG TAA AGA TTC GGA AGG GCA GCC AGC AAC AGG CCA CCC TGC AAT GGC ATA TTG CAT GGT GTG CTC CTT ATT TAT ACA TAA CGA AAA ACG CCT CGA GTG AAG CGT TAT TGG TAT GCG GTA AAA CCG CAC TCA GGC GGC CTT GAT AGT CAT ATC ATC TGA ATC AAA TAT TCC TGA TGT ATC GAT ATC GGT AAT TCT TAT TCC TTC GCT ACC ATC CAT TGG AGG CCA TCC TTC CTG ACC ATT TCC ATC ATT CCA GTC GAA CTC ACA CAC AAC ACC ATA TGC ATT TAA GTC GCT TGA AAT TGC TAT AAG CAG AGC ATG TTG CGC CAG CAT GAT TAA TAC AGC ATT TAA TAC AGA GCC GTG TTT ATT GAG TCG GTA TTC AGA GTC TGA CCA GAA ATT ATT AAT CTG GTG AAG TTT TTC CTC TGT CAT TAC GTC ATG GTC GAT TTC AAT TTC TAT TGA TGC TTT CCA GTC GTA ATC AAT GAT GTA TTT TTT GAT GTT TGA CAT CTG TTC ATA TCC TCA CAG ATA AAA AAT CGC CCT CAC ACT GGA GGG CAA AGA AGA TTT CCA ATA ATC |

| | |
|---|---|
| 400 bp DNA with 1× p21 binding motif for *in vitro* droplet assay | CTT TTG CTT GAT CTC AGT TTC AGT ATT AAT ATC CAT TTT<br>TTA TAA GCG TCG ACG GCT TCA CGA AAC ATC TTT TCA TCG<br>CCA ATA AAA GTG GCG ATA GTG AAT TTA GTC TGG ATA GCC<br>ATA AGT GTT TGA TCC ATT CTT TGG GAC TCC TGG CTG ATT<br>AAG TAT GTC GAT AAG GCG TTT CCA TCC GTC ACG TAA TTT<br>ACG GGT GAT TCG TTC AAG TAA AGA TTC GGA AGG GCA<br>GCC AGC AAC AGG CCA CCC TGC AAT GGC ATA TTG CAT<br>GGT GTG CTC CTT ATT TAT ACA TAA CGA AAA ACG CCT CGA<br>GTG AAG CGT TAT TGG TAT GCG GTA AAA CCG CAC TCA<br>GGC GGC CTT GAT AGT CAT ATC ATC TGA ATC AAA CAA CAT<br>GTT GGG ACA TGT TCT CAC T |
| 400 bp DNA with 2× p21 binding motifs for *in vitro* droplet assay | AGT GAG AAC ATG TCC CAA CAT GTT GTT AAT ATC CAT TTT<br>TTA TAA GCG TCG ACG GCT TCA CGA AAC ATC TTT TCA TCG<br>CCA ATA AAA GTG GCG ATA GTG AAT TTA GTC TGG ATA GCC<br>ATA AGT GTT TGA TCC ATT CTT TGG GAC TCC TGG CTG ATT<br>AAG TAT GTC GAT AAG GCG TTT CCA TCC GTC ACG TAA TTT<br>ACG GGT GAT TCG TTC AAG TAA AGA TTC GGA AGG GCA<br>GCC AGC AAC AGG CCA CCC TGC AAT GGC ATA TTG CAT<br>GGT GTG CTC CTT ATT TAT ACA TAA CGA AAA ACG CCT CGA<br>GTG AAG CGT TAT TGG TAT GCG GTA AAA CCG CAC TCA<br>GGC GGC CTT GAT AGT CAT ATC ATC TGA ATC AAA CAA CAT<br>GTT GGG ACA TGT TCT CAC T |
| 400 bp DNA with 3× p21 binding motifs for *in vitro* droplet assay | AGT GAG AAC ATG TCC CAA CAT GTT GTT AAT ATC CAT TTT<br>TTA TAA GCG TCG ACG GCT TCA CGA AAC ATC TTT TCA TCG<br>CCA ATA AAA GTG GCG ATA GTG AAT TTA GTC TGG ATA GCC |

| | |
|---|---|
| | ATA AGT GTT TGA TCC ATT CTT TGG GAC TCC TGG CTG ATT AAG TAT GTC GAT AAG GCG TTT CCA TCC GTC ACG TAA TTT ACG GGT GAT TCG TTC AAG TAA AGA TTC GGA AGG GCA GCC AGC AAC AGG CCA CCC TGC AAT GGC ATA TTG CAT GGT GTG CTC CTT ATT TAT ACA TAA CGA AAA ACG CCT CGA GTG AAG CGT TAT TGG TAT GCG GTA AAA CAA CAT GTT GGG ACA TGT TCT CTT CTA TGC CAG AGC TCA ACA TGT TGG GAC ATG TTC CTC GAG CGG T |
| 400 bp with 4× p21 binding motifs for *in vitro* droplet assay | ACC GCT CGA GGA ACA TGT CCC AAC ATG TTG AGC TCT GGC ATA GAA GAG AAC ATG TCC CAA CAT GTT GTC TTT TCA TCG CCA ATA AAA GTG GCG ATA GTG AAT TTA GTC TGG ATA GCC ATA AGT GTT TGA TCC ATT CTT TGG GAC TCC TGG CTG ATT AAG TAT GTC GAT AAG GCG TTT CCA TCC GTC ACG TAA TTT ACG GGT GAT TCG TTC AAG TAA AGA TTC GGA AGG GCA GCC AGC AAC AGG CCA CCC TGC AAT GGC ATA TTG CAT GGT GTG CTC CTT ATT TAT ACA TAA CGA AAA ACG CCT CGA GTG AAG CGT TAT TGG TAT GCG GTA AAA CAA CAT GTT GGG ACA TGT TCT CTT CTA TGC CAG AGC TCA ACA TGT TGG GAC ATG TTC CTC GAG CGG T |

#### 3. Protein sequences

| Protein | Sequence (TAD, PRD, DBD, HD, OD and CTD) |
|---|---|
| 6×His-3×Flag-p53$^{4M}$ | MGSSHHHHHHAMAMEEPQSDPSVEPPLSQETFSDLWKLLPENNVLSPLPSQAMDDLMLSPDDIEQWFTEDPGPDEAPRMPEAAPRVAPAPAAPTPAAPAPAPSWPLSSSVPSQKTYQGSYGFRLGFLHSGTAKSVTCTYSPALNKLFCQLAKTCPVQLWVDSTPPPGTRVRAMAIYKQSQHMTEVVRRCPHHERCSDSDGLAPPQHLIRVEGNLRAEYLDDRNTFRHSVVVPYEPPEVGSDCTTIHYNYMCYSSCMGGMNRRPILTIITLEDSSGNLLGRDSFEVRVCACAGRDRRTEEENLRKKGEPHHELPPGSTKRALPNNTSSSPQPKKKPLDGEYFTLQIRGRERFEMFRELNEALELKDAQAGKEPGGSRAHSSHLKSKKGQSTSRHKKLMFKTEGPDSD* |
| 6×His-PRD-DBD-TET-CTD (p53$^{4M}$ ΔTAD) | MGSSHHHHHHAMAAPRMPEAAPRVAPAPAAPTPAAPAPAPSWPLSSSVPSQKTYQGSYGFRLGFLHSGTAKSVTCTYSPALNKLFCQLAKTCPVQLWVDSTPPPGTRVRAMAIYKQSQHMTEVVRRCPHHERCSDSDGLAPPQHLIRVEGNLRAEYLDDRNTFRHSVVVPYEPPEVGSDCTTIHYNYMCYSSCMGGMNRRPILTIITLEDSSGNLLGRDSFEVRVCACAGRDRRTEEENLRKKGEPHHELPPGSTKRALPNNTSSSPQPKKKPLDGEYFTLQIRGRERFEMFRELNEALELKDAQAGKEPGGSRAHSSHLKSKKGQSTSRHKKLMFKTEGPDSD* |

### 4. Supplementary Figures

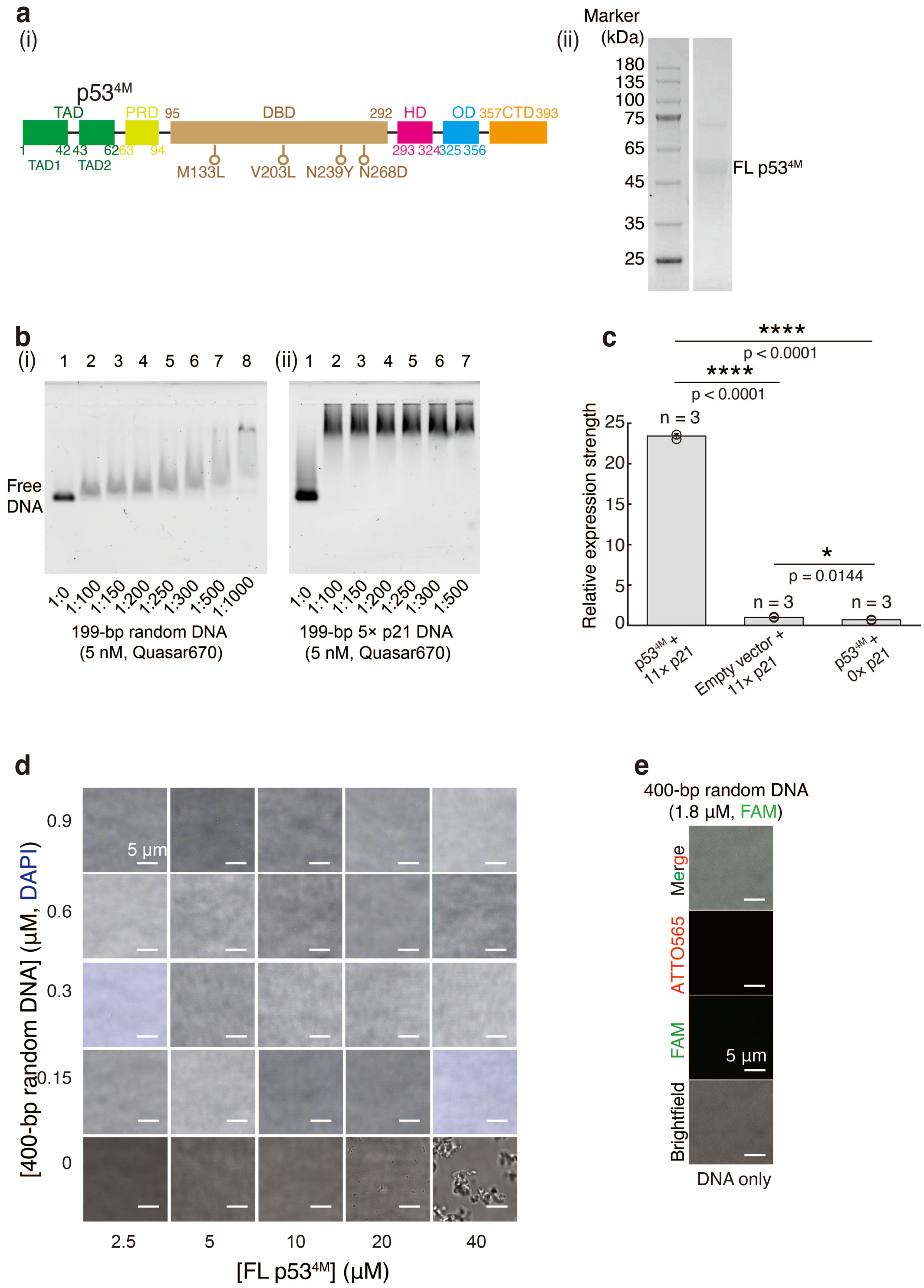


**Supplementary Fig. 1. Full length p53$^{4M}$ cannot form the DPICs with random DNA.**

(**a**) Schematic representation and SDS-PAGE analysis of p53$^{4M}$. (**b**) EMSAs (1% agarose gel) showing $p53^{4M}$ interaction with 199-bp DNA. (i) 199-bp random DNA; (ii) 199-bp DNA containing 5× p21 binding motifs. DNA substrates were labeled with Quasar670 and imaged using an Amersham Typhoon RGB system (with a 635 nm laser and Cy5 670BP30 filter). SDS-PAGE in a(ii) and EMSAs in b were conducted three times (n = 3). (**c**) Luciferase assays using supernatants from H1299 cells transiently transfected with $p53^{4M}$-expressing vector or empty vector, and a firefly luciferase vector containing 0× or 11× p21 promoter sequences preceding a minimal promoter. Data were normalized to the luciferase activity of the empty vector control. Three independent biological replicates were performed for each condition (n = 3). Error bars represent mean ± s.d. Statistical significances were evaluated based on the student's t-test. P value: two-tailed; p value style: GP: 0.1234 (ns), 0.0332 (*), 0.0021 (**), 0.0002 (***), <0.0001 (****). Confidence level: 95%. (**d**) *In vitro* droplet experiments by mixing 0, 0.15, 0.3, 0.6, and 0.9 μM 400-bp random DNA with 2.5, 5, 10, 20, and 40 μM dark $p53^{4M}$. DNA was labeled with DAPI. (**e**) *In vitro* droplet experiment using 1.8 μM 400-bp random DNA labeled with FAM. *In vitro* droplet experiments were conducted in a working buffer of 8 mM Tris-HCl (pH 7.5), 120 mM NaCl, 4% glycerol, and 16 mM DTT, without any crowding agents. Independent *in vitro* droplet experiments in d and e were repeated three times (n = 3) each.

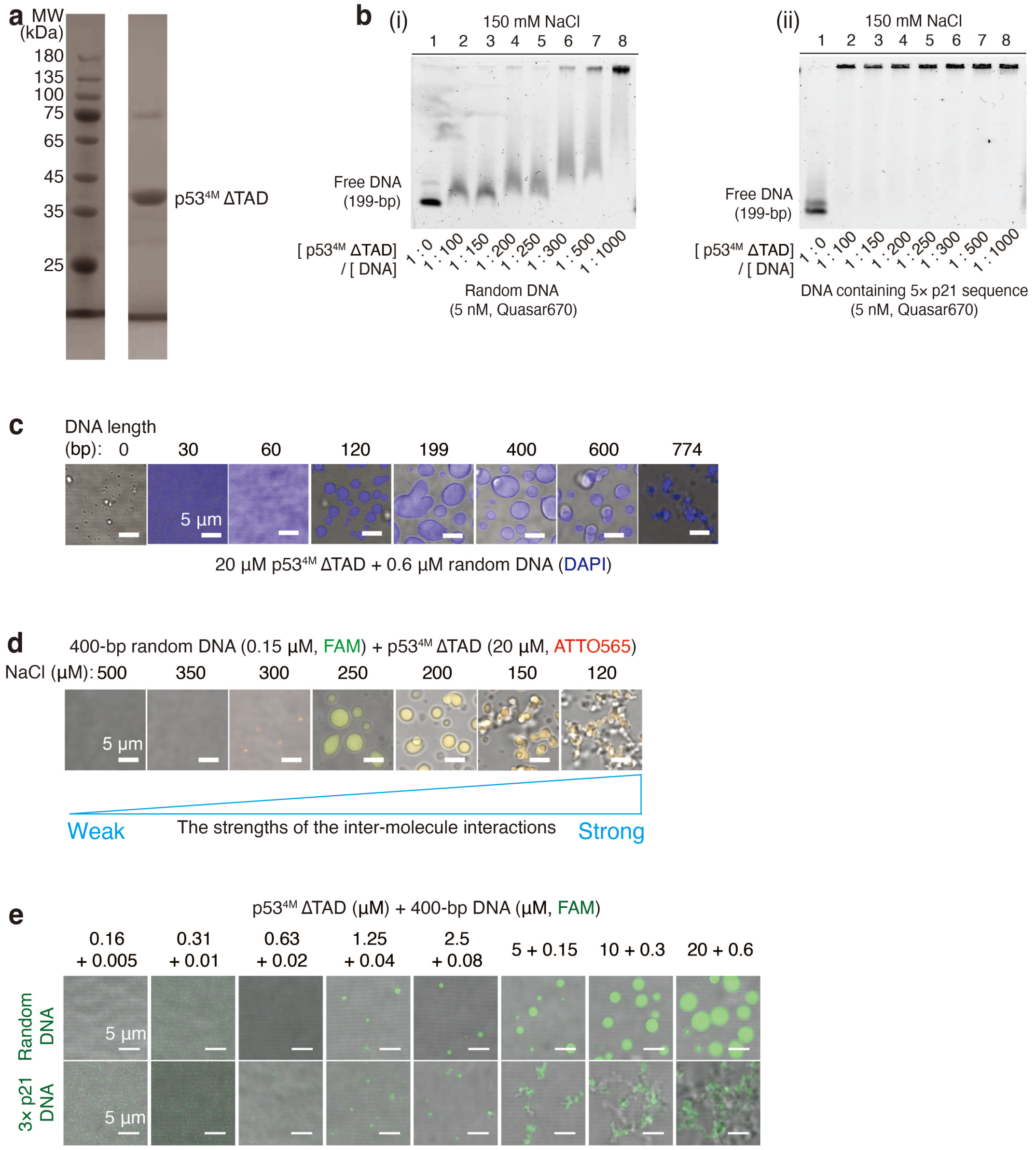


**Supplementary Fig. 2. Purification of *in vitro* p53$^{4M}$ ΔTAD.** (**a**) SDS-PAGE analysis of p53$^{4M}$ ΔTAD. (**b**) EMSAs (1% agarose gel) illustrating the interaction of p53$^{4M}$ ΔTAD with 199-bp DNA. (i) 199-bp random DNA; (ii) 199-bp DNA containing 5× p21 binding motifs. DNA substrates were labeled with Quasar670 and imaged using an Amersham Typhoon RGB system (with a 635 nm laser and Cy5 670BP30 filter). (**c**) *In vitro* droplet experiments by combining 20 µM p53$^{4M}$ ΔTAD and 0.6 µM 0, 30, 60, 120, 199, 400, 600, and 774-bp random DNA labeled with DAPI. (**d**) *In vitro* droplet experiments by mixing 20 µM p53$^{4M}$

ΔTAD labeled with ATTO565 and 0.15 μM 400-bp DNA labeled with FAM at different NaCl concentrations: 500, 350, 300, 250, 200, 150, and 120 mM. Except for the NaCl concentration, the other components of the working buffer were consistent with those used in other droplet experiments. (**e**) *In vitro* droplet experiments by mixing $p53^{4M}$ ΔTAD and 400-bp DNA labeled with FAM at varying concentrations. The working buffer for *in vitro* droplet experiments was 8 mM Tris-HCl (pH 7.5), 120 mM NaCl, 4% Glycerol, and 16 mM DTT. The working buffer contained no crowding agents in this study. All experiments were repeated three times (n = 3).

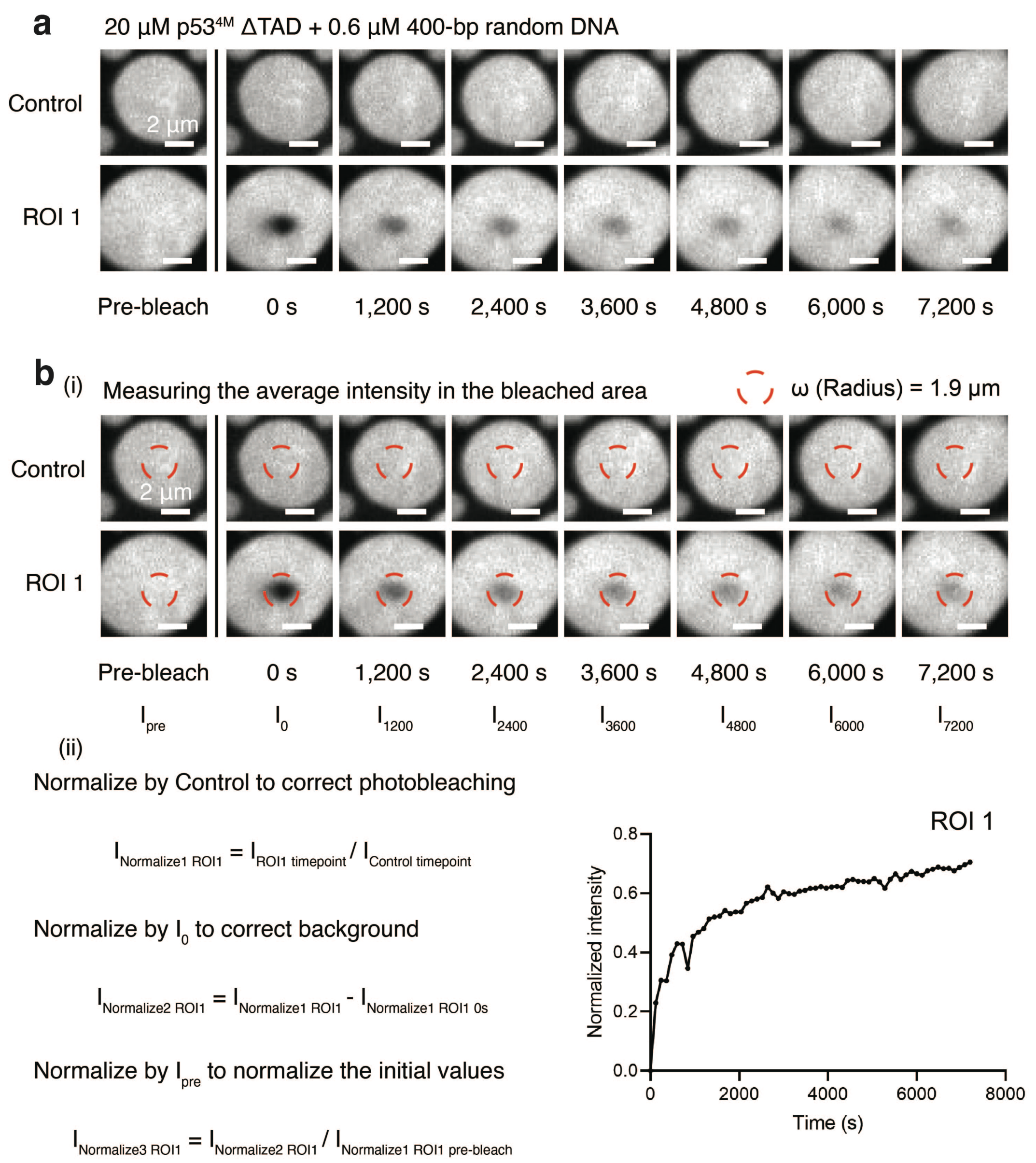


**Supplementary Fig. 3. Data analysis for the FRAP experiment.** (**a**) FRAP experiment conducted on the droplet-like DPIC composed of 20 μM p53$^{4M}$ ΔTAD and 0.6 μM 400-bp random DNA. p53$^{4M}$ ΔTAD was labeled with ATTO565, and DNA was labeled with FAM. Eight panels display the time course of the selected droplet, with time indicated at the bottom in seconds. “Control” denotes the droplet without photobleaching, while “ROI 1” corresponds to the droplet subjected to photobleaching. ROI stands for the region of interest. (**b**) The intensity within the red dashed circle (i) was utilized for data analysis (ii).

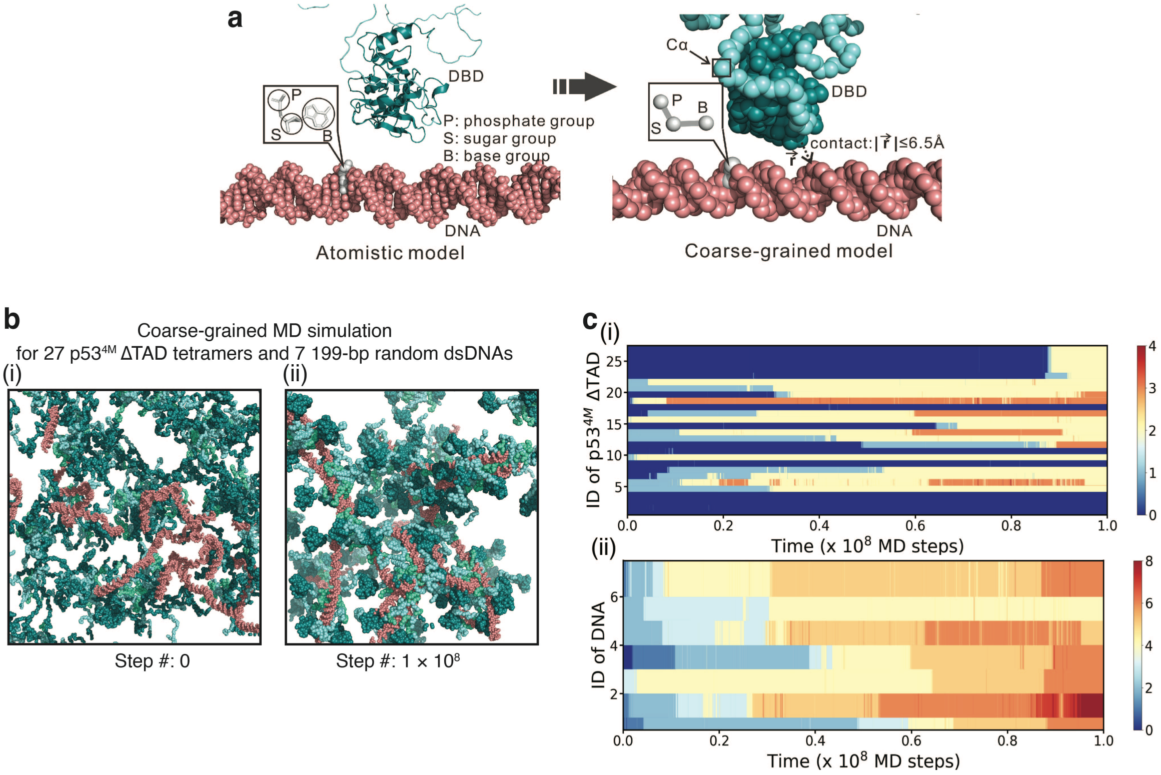


**Supplementary Fig. 4. Coarse-grained MD simulations for DPIC formation.** (**a**) Illustration of the coarse-grained model depicting the DBD domain of p53$^{4M}$ and a random DNA molecule. Residues are represented by a single particle at their $C_{\alpha}$ position, while nucleotides are depicted by three beads corresponding to phosphate, sugar, and base. (**b**) Three-dimensional structures for the initial configuration (i) and the final configuration of MD simulations (ii) in a system containing 27 p53$^{4M}$ ΔTAD tetramers and 7 199-bp dsDNAs at a temperature of 300 K and a salt concentration of 150 mM. (**c**) (i) Number of bound dsDNA for each p53$^{4M}$ ΔTAD tetramer over time in an MD trajectory. (ii) Number of bound p53$^{4M}$ ΔTAD tetramers for each dsDNA chain over time. Color bars represent the quantity of protein (DNA) bound to a specific DNA (protein). For assigning the bound state, contact between residue beads and nucleotide beads was considered formed if their distance was less than 6.5 Å (a).

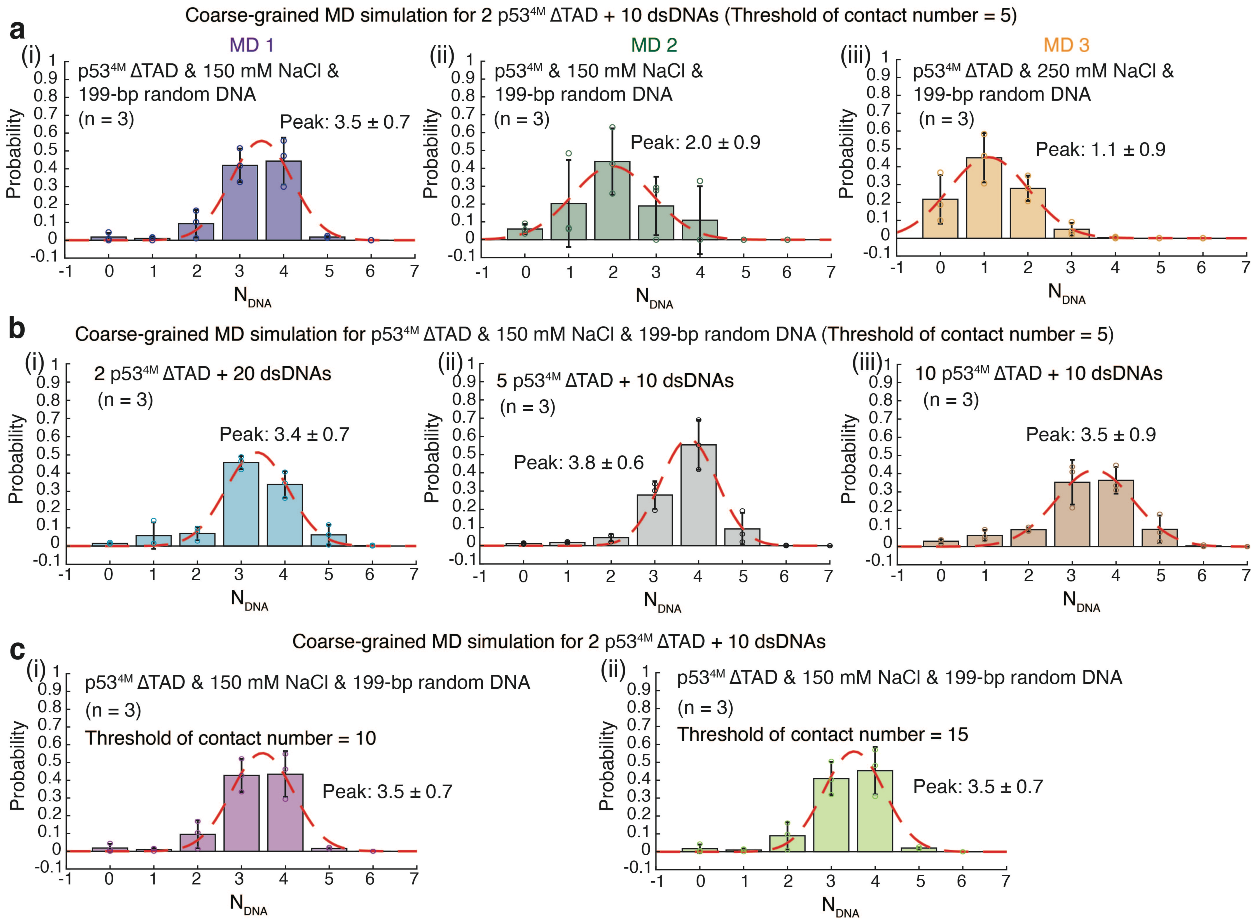


**Supplementary Fig. 5. Capacity of p53$^{4M}$ ΔTAD tetramer to bridge multiple dsDNA chains.** (**a**) MD simulations were performed in a smaller system comprising 2 p53$^{4M}$ ΔTAD tetramers and 10 dsDNAs. The contact number threshold between p53$^{4M}$ monomer and dsDNA was set to 5 (Threshold of contact number = 5). (i) Probability distribution of the number of bound 199-bp random dsDNA for each p53$^{4M}$ ΔTAD tetramer (MD 1: blue) at 300 K and a salt concentration of 150 mM; (ii) Probability distribution of the number of bound 199-bp random dsDNA for each full-length p53$^{4M}$ tetramer (MD 2: green) at 300 K and a salt concentration of 150 mM; (iii) Probability distribution of the number of bound 199-bp random dsDNA for each p53$^{4M}$ ΔTAD tetramer (MD 3: orange) at 300 K and a salt concentration of 250 mM. (**b**) Probability distribution of the number of bound 199-bp random dsDNA for each p53$^{4M}$ ΔTAD tetramer at 300 K and a salt concentration of 150 mM (Threshold of contact number = 5). (i) A smaller system containing 2 p53$^{4M}$ ΔTAD tetramers and 20 dsDNAs was used for MD simulations; (ii) A smaller system containing 5 p53$^{4M}$ ΔTAD tetramers and 10 dsDNAs was used for MD simulations; (iii) A smaller system containing 10 p53$^{4M}$ ΔTAD tetramers and 10 dsDNAs was used for MD simulations. (**c**) A smaller system containing 2 p53$^{4M}$ ΔTAD tetramers and 10 random dsDNAs was used for MD simulations. Probability distribution of the number of bound 199-bp dsDNA for each p53$^{4M}$ ΔTAD tetramer at 300 K and a salt concentration of 150 mM: (i) Threshold of contact number = 10; (ii) Threshold of contact number = 15. Probabilities were calculated based on snapshots from three independent MD simulations (n = 3). Snapshots of the first $3\times 10^7$ MD steps were omitted in calculating the probabilities. Error bars represent mean ± s.d. All probability distributions in a-d were fitted with a 1D Gaussian function (red dash line). Error bars of the fitting parameters represent 95% confidence intervals obtained through Gaussian function fitting.

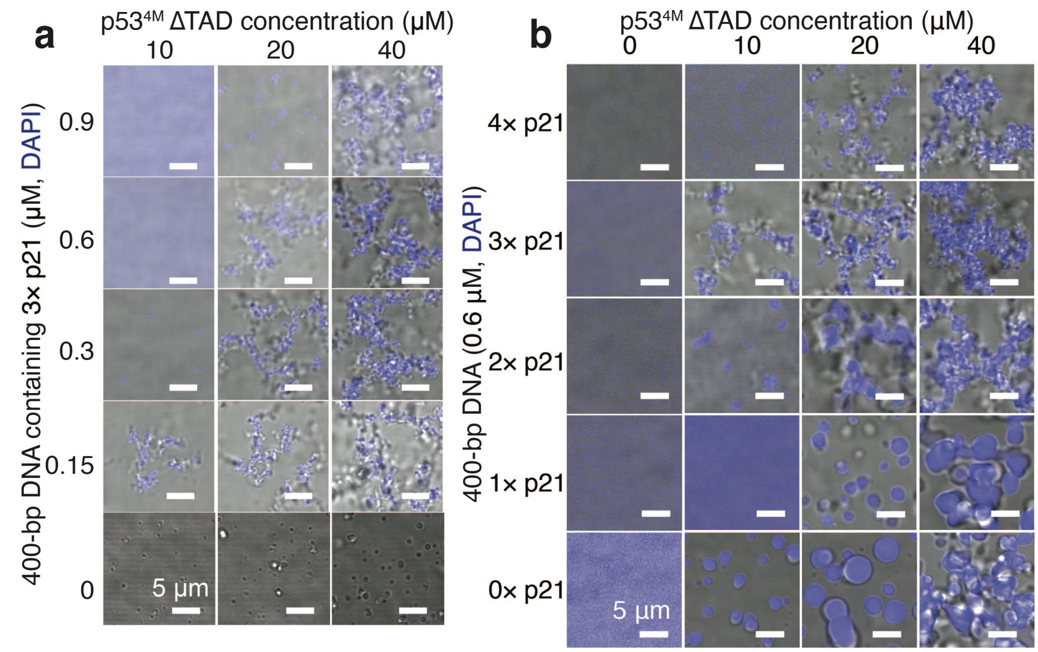


**Supplementary Fig. 6. Morphological transition of DPICs from droplet-like to "pearl chain"-like induced by the addition of p21 binding motifs into the DNA substrates.** (**a**) *In vitro* droplet experiments involving the mixing of 0, 0.15, 0.3, 0.6, and 0.9 μM 400-bp DNA containing 3× p21 binding motifs with 10, 20, and 40 μM dark p53$^{4M}$ ΔTAD. (**b**) *In vitro* droplet experiments by mixing 0.6 μM 400-bp DNA containing 0-4× p21 binding motifs with 0, 10, 20, and 40 μM dark p53$^{4M}$ ΔTAD.

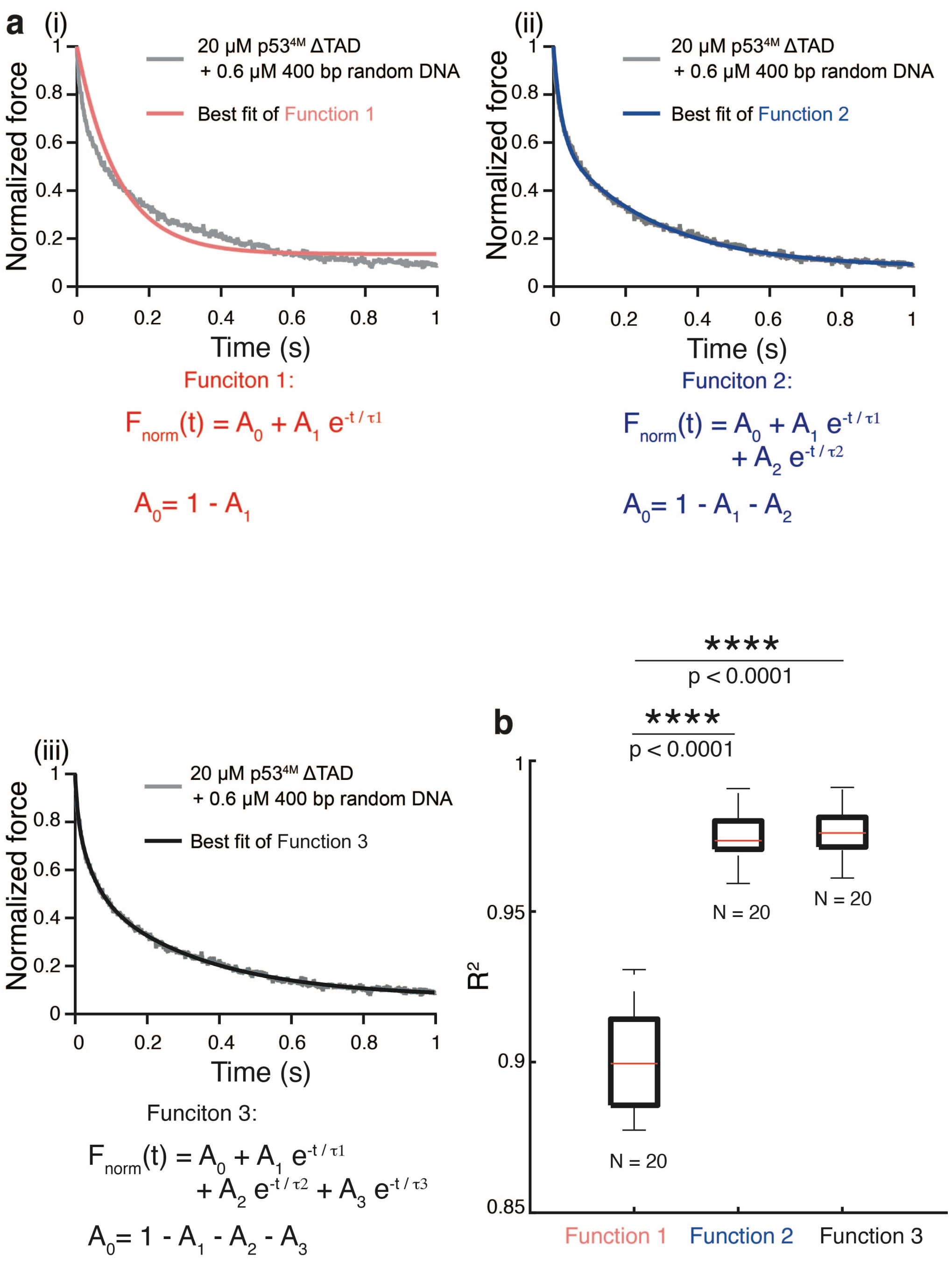


**Supplementary Fig. 7. Discussion on the Generalized Maxwell Model.** (**a**) Representative normalized relaxation curves of DPICs formed by 20 μM p53$^{4M}$ ΔTAD and 0.6 μM 400-bp random DNA (gray). The pink line (i), blue line (ii), and black line (iii) represent theoretical fitting curves by three generalized Maxwell models: Function 1 (i), Function 2 (ii), and Function 3 (iii). (**b**) Boxplot of $R^2$ for the three fitting functions. The total number (N) of DPICs examined in a single AFM-FS experiment: N = 20 for the condition of 20 μM p53$^{4M}$ ΔTAD and 0.6 μM 400-bp random DNA. For the boxplot, the red bar represents median. The bottom edge of the box represents 25$^{th}$ percentiles, and the top is 75$^{th}$ percentiles. Most extreme data points are covered by the whiskers except outliers. The '+' symbol is used to represent the outliers. Statistical significance was analyzed using unpaired t test for two groups. P value: two-tailed; p value style: GP: 0.1234 (ns), 0.0332 (*), 0.0021 (**), 0.0002 (***), <0.0001 (****). Confidence level: 95%.

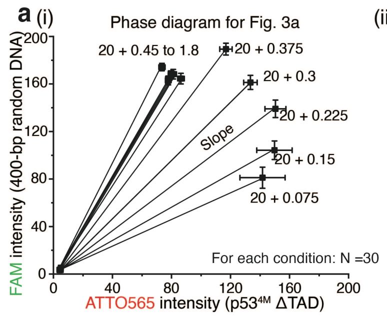


(ii)

| Initial concentration of protein (μM) | Initial concentration of DNA (μM) | Slope in phase diagram |
|---|---|---|
| 20 | 1.8 | 2.2 |
| 20 | 1.5 | 2.2 |
| 20 | 1.2 | 2.5 |
| 20 | 0.9 | 2.2 |
| 20 | 0.75 | 2.2 |
| 20 | 0.6 | 2.2 |
| 20 | 0.45 | 2.0 |
| 20 | 0.375 | 1.7 |
| 20 | 0.3 | 1.2 |
| 20 | 0.225 | 0.9 |
| 20 | 0.15 | 0.7 |
| 20 | 0.075 | 0.6 |

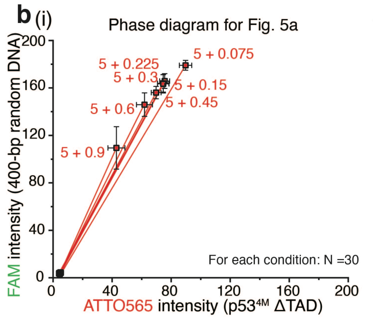


(ii)

| Initial concentration of protein (μM) | Initial concentration of DNA (μM) | Slope in phase diagram |
|---|---|---|
| 5 | 0.9 | 2.8 |
| 5 | 0.6 | 2.5 |
| 5 | 0.45 | 2.3 |
| 5 | 0.3 | 2.3 |
| 5 | 0.225 | 2.3 |
| 5 | 0.15 | 2.3 |
| 5 | 0.075 | 2.1 |

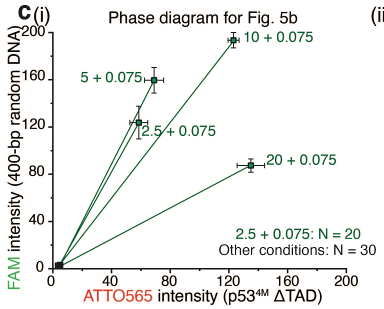


(ii)

| Initial concentration of protein (μM) | Initial concentration of DNA (μM) | Slope in phase diagram |
|---|---|---|
| 20 | 0.075 | 0.6 |
| 10 | 0.075 | 1.6 |
| 5 | 0.075 | 2.4 |
| 2.5 | 0.075 | 2.3 |

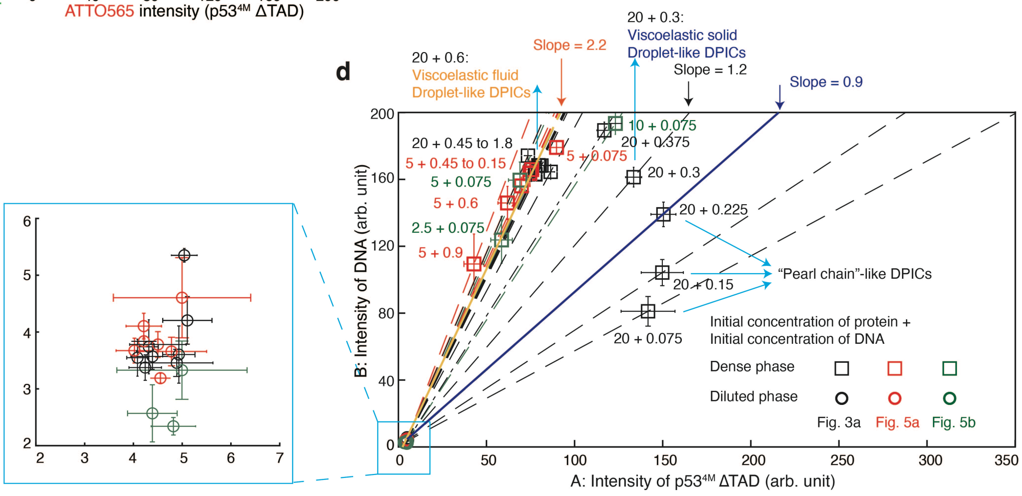


**Supplementary Fig. 8. Tie lines of DPICs formed by the combination of p53$^{4M}$ ΔTAD and 400-bp random DNA.** (**a-c**) (i) Two-component phase diagram of DPICs derived from three sets of two-color *in vitro* droplet assays. (ii) Slopes of the tie lines in a. Data extracted from Fig. 3a (a), data from Fig. 5a (b), and Data from Fig. 5b (c). (**d)** Two-component phase diagram of DPIC from all three data sets in a-c. The intensity of p53$^{4M}$ ΔTAD (ATTO565) and 400-bp random DNA (FAM) in both the dense (square) and dilute phases (circle) from all conditions in Fig. 3a (black), 5a (red), and 5b (green) was plotted. The box zooms in on all data points in the dilute phase. The numbers labeling the dense or dilute phase represent the initial concentration of p53$^{4M}$ ΔTAD and the initial concentration of 400-bp random DNA. The dashed line connecting data points of the dense phase and dilute phase for each condition represents the tie line. Two slope thresholds of tie lines are highlighted: (i) the orange line (slope ~2.2), representing a phase transition between viscoelastic fluid and viscoelastic solid; (ii) the blue line (slope ~0.9), representing a morphology transition between DPICs with droplet-like morphology and "pearl chain"-like structure.

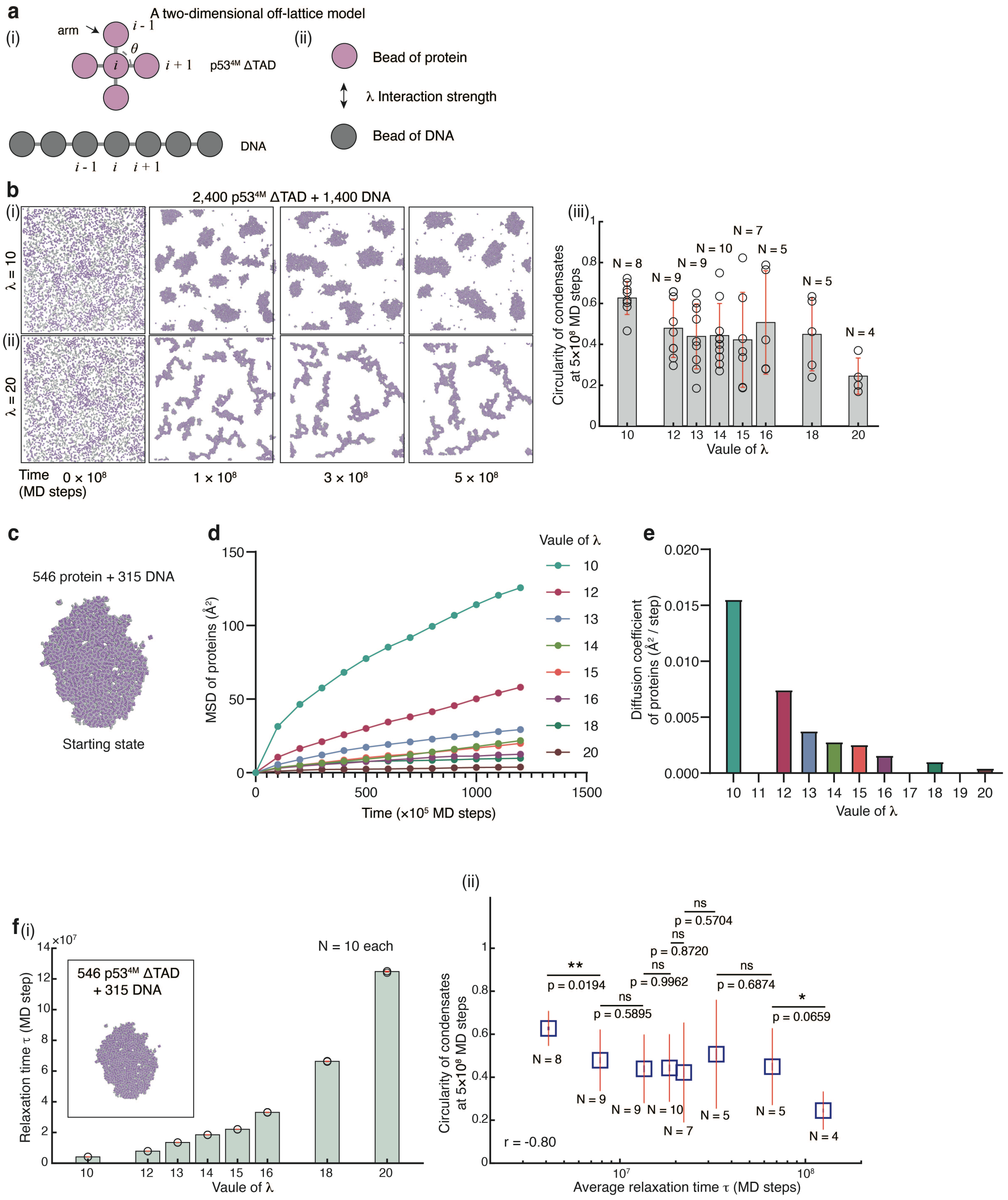


**Supplementary Fig. 9. Interplay between relaxation time and morphology: simulations with a two-dimensional off-lattice model.** (**a**) Schematic illustration of the two-dimensional off-lattice model. (i) Simplified structure of p53 tetramer and dsDNA. (ii) The interaction strength between the branching bead of p53$^{4M}$ ΔTAD tetramer and the bead on DNA was controlled by a parameter λ. (**b**) DPIC formation in the simulation system containing 2,400 $p53^{4M}$ ΔTAD tetramers and 1,400 DNA chains with $\lambda$ = 10 (i) or $\lambda$ = 20 (ii). The circularity of DPICs formed after $5\times 10^8$ steps of simulations with different $\lambda$ was shown in (iii). The total number of DPICs (N) formed in each simulation and used for the calculation of the circularity was listed. (**c**) Initial configuration of the condensate comprising 546 $p53^{4M}$ ΔTAD and 315 DNA chains. (**d**) Mean Squared Displacement (MSD) of proteins within the condensate in a with varying values of $\lambda$ during 0 to $1{,}250\times 10^5$ MD steps. (**e**) Diffusion coefficient values of proteins obtained from the MSD in d. (**f**) Dynamic property of DPICs with different $\lambda$. (i) The value of relaxation time calculated from the diffusion coefficient in c-e. We used 10 snapshots from the diffusion simulations to extract the droplet radius in estimating the relaxation time. The box shows the starting structure of the condensate containing 546 $p53^{4M}$ ΔTAD and 315 DNA chains. (ii) Correlation between estimated relaxation time and DPIC circularity given in b(iii) and f(i). Error bars in b and f represent mean ± s.d. Statistical significance was analyzed using unpaired t test for two groups. P value: two-tailed; p value style: GP: 0.1234 (ns), 0.0332 (*), 0.0021 (**), 0.0002 (***), <0.0001 (****). Confidence level: 95%.

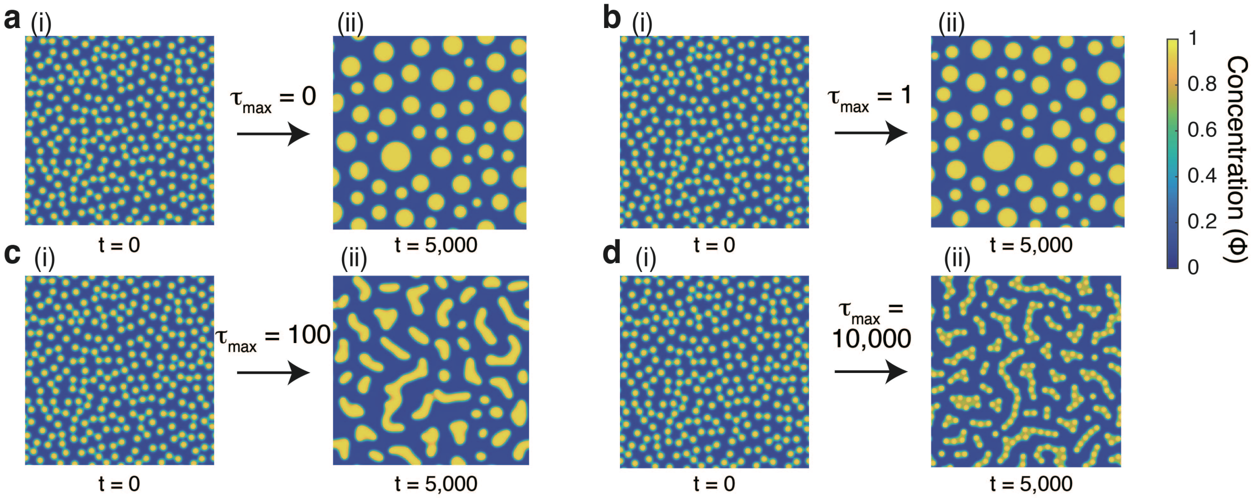


**Supplementary Fig. 10. Interplay between relaxation time and morphology: simulations with a mathematical model based on VPS theory.** (**a-d**) Results by a mathematical model based on the VPS theory. Droplets with the same size were randomly placed in a square box at the initial state (a(i), b(i), c(i), and d(i)), then underwent coalescence after 5,000 steps with $\tau_{max}$ = 0 a(ii), $\tau_{max}$ = 1 b(ii), $\tau_{max}$ = 100 c(ii), and $\tau_{max}$ = 10,000 d(ii). The color represented the concentration of substrates ($\phi$).

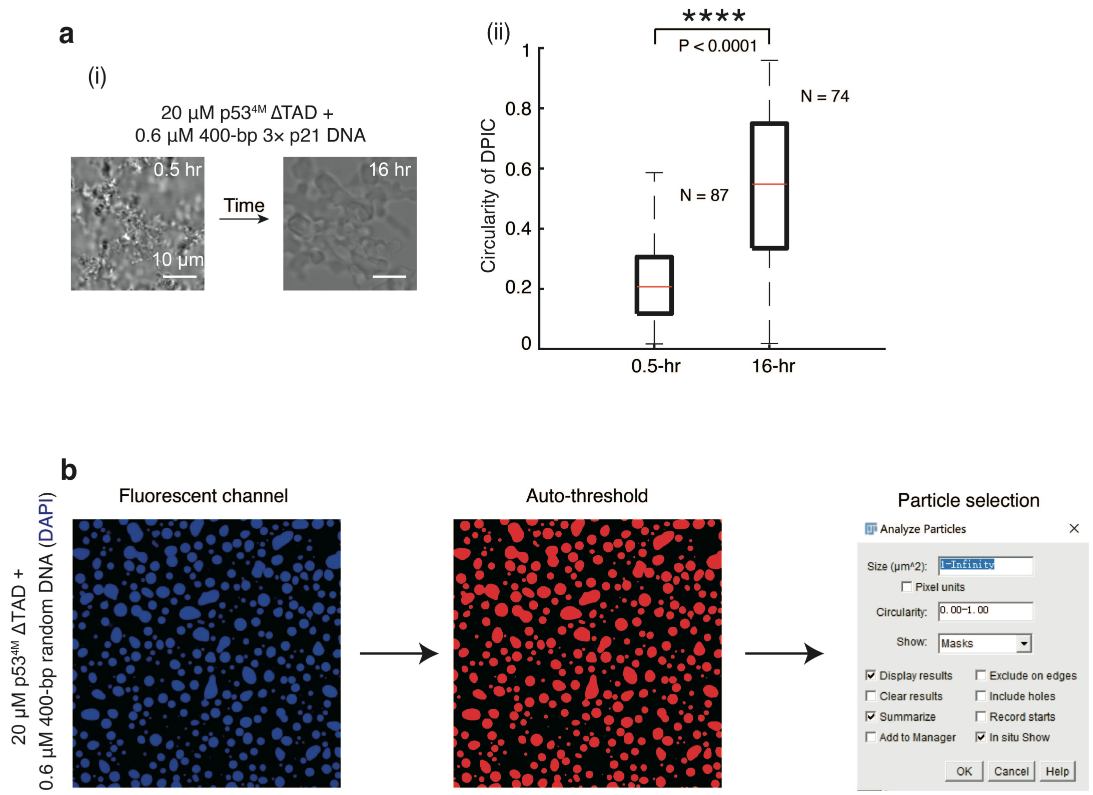


**Supplementary Fig. 11. Dynamic behaviors of "Pearl chain"-like DPICs.** (**a**) (i) Time course panels of DPIC formed by 20 μM p53$^{4M}$ ΔTAD and 0.6 μM 400-bp DNA containing 3× p21 binding motifs in one *in vitro* droplet experiment, with time in hours. (ii) Boxplot depicting DPIC circularity for the 0.5-hour and 16-hour conditions in (i). The total number (N) of DPICs examined over a single *in vitro* droplet experiment in a(ii): N = 87 for the 0.5-hour condition and N = 74 for the 16-hour condition. For the boxplot, the red bar represents median. The bottom edge of the box represents 25$^{th}$ percentiles, and the top is 75$^{th}$ percentiles. Most extreme data points are covered by the whiskers except outliers. The '+' symbol is used to represent the outliers. Statistical significance was analyzed using unpaired t test for two groups. P value: two-tailed; p value style: GP: 0.1234 (ns), 0.0332 (*), 0.0021 (**), 0.0002 (***), <0.0001 (****). Confidence level: 95%. (**b**) Droplet circularity analysis for *in vitro* droplet assay in a. Particle selection and circularity calculation: 20 μM p53$^{4M}$ ΔTAD and 0.6 μM random DNA, and DNA was labeled with DAPI. The subroutine of "analyze particles" in Fiji (Analyze/Analyze particles) was used to track the image events (**Methods**).

### 5. Supplementary Movie Legends

**Supplementary Movie 1. 20 μM p53$^{4M}$ ΔTAD and 0.6 μM 400-bp random DNA formed the droplet-like DPICs.** The video was acquired for 4 hours after a 30-min incubation. 4-min shutter time was used.

**Supplementary Movie 2. Coarse-grained MD simulation for the system containing 7 199-bp random dsDNA chains and 27 p53$^{4M}$ ΔTAD tetramers at the temperature of 300 K and the salt concentration of 150 mM.** The video corresponds to the initial stage of the full MD trajectory. The time interval between the frames was $1 \times 10^5$ MD steps. One can observe the development of the DNA-protein bridging interactions with the time evolution.

**Supplementary Movie 3. Coarse-grained MD simulation for the system containing 10 199-bp random dsDNA chains and 2 p53$^{4M}$ ΔTAD tetramers at the temperature of 300 K and the salt concentration of 150 mM.** For a better view, only one p53$^{4M}$ tetramer was focused in the video. The time interval between the frames was $1 \times 10^5$ MD steps. One can observe that one p53$^{4M}$ ΔTAD tetramer can capture multiple DNA chains simultaneously during the simulation.

**Supplementary Movie 4. Coarse-grained MD simulation for the system containing 10 199-bp random dsDNA chains and 2 full-length $p53^{4M}$ tetramers at the temperature of 300 K and the salt concentration of 150 mM.** For a better view, only one $p53^{4M}$ tetramer was focused in the video. One can observe that the number of DNA chains engaged by one full-length $p53^{4M}$ tetramer becomes fewer compared to the case of the $p53^{4M}$ ΔTAD tetramer.

**Supplementary Movie 5. Coarse-grained MD simulation for the system containing 10 199-bp random dsDNA chains and 2 $p53^{4M}$ ΔTAD tetramers at the temperature of 300 K and the salt concentration of 250 mM.** For a better view, only one $p53^{4M}$ tetramer was focused in the video. One can observe that the number of DNA chains engaged by one $p53^{4M}$ tetramer becomes fewer due to the increasing of the salt concentration.

**Supplementary Movie 6. 20 μM $p53^{4M}$ ΔTAD and 0.6 μM 400-bp DNA containing 3× p21 binding motifs formed the granular DPICs.** The video was acquired for 4 hours after a 30-min incubation. 4-min shutter time was used.

**Supplementary Movie 7. Representative simulation trajectory of a two-dimensional off-lattice model containing 2,400 protein molecules and 1,400 DNA chains with $\lambda = 10$.** The temperature of the simulation is 298 K. The simulation lasted for a total of $5 \times 10^8$ MD steps.

**Supplementary Movie 8. Representative simulation trajectory of a two-dimensional off-lattice model containing 2,400 protein molecules and 1,400 DNA chains with λ = 20.** The temperature of the simulation is 298 K. The simulation lasted for a total of 5 × $10^8$ MD steps.

**Supplementary Movie 9. Numerical simulation of viscoelastic phase separation based on the two-fluid model with $\tau_{max}$ = 0.** In the numerical simulation, parameters were set: $\chi = 3$, $G_B = G_S = 10$ and $\phi_c = 0.7$. The maximum relaxation times ($\tau_{max}$) of $G_B$ and $G_S$ were set as 0. At the initial, 265 condensates with the initial radius ($R_{\text{ini}}$) as 4 were placed randomly in a 2D 511*511 grid with the grid size ($\Delta l$) as 0.25. The total time ($t_{\text{tot}}$) for the simulation is 5000 and the time interval ($\Delta t$) for the simulation is 0.005. One can observe the droplets undergo coalescence.

**Supplementary Movie 10. Numerical simulation of viscoelastic phase separation based on the two-fluid model with $\tau_{max}$ = 1.** All other parameters are the same to Supplementary Movie 11, except the maximum relaxation times ($\tau_{max}$) of $G_B$ and $G_S$ were set as 1.

**Supplementary Movie 11. Numerical simulation of viscoelastic phase separation based on the two-fluid model with $\tau_{max}$ = 100.** All other parameters are the same to Supplementary Movie 11, except the maximum relaxation times ($\tau_{max}$) of $G_B$ and $G_S$ were set as 100.

**Supplementary Movie 12. Numerical simulation of viscoelastic phase separation based on the two-fluid model with $\tau_{max}$ = 10,000.** All other parameters are the same to Supplementary Movie 11, except the maximum relaxation times ($\tau_{max}$) of $G_B$ and $G_S$ were set as 10,000.
